# Solid tumor-on-chip model for efficacy and safety assessment of CAR-T cell therapy

**DOI:** 10.1101/2023.07.13.548856

**Authors:** Tengku Ibrahim Maulana, Claudia Teufel, Madalena Cipriano, Lisa Lazarevski, Francijna E. van den Hil, Valeria Orlova, André Koch, Miriam Alb, Michael Hudecek, Peter Loskill

**Affiliations:** Department of Microphysiological Systems, Institute of Biomedical Engineering, Faculty of Medicine, Eberhard-Karls-University-Tübingen, Tübingen, Germany; NMI Natural and Medical Sciences Institute at the University of Tübingen, Reutlingen, Germany; 3R Center Tübingen for In Vitro Models and Alternatives to Animal Testing, Tübingen, Germany; Department of Anatomy and Embryology, Leiden University Medical Center, Leiden, The Netherlands; Department of Women’s Health Tübingen, Eberhard-Karls-University Tübingen, Tübingen, Germany; Medizinische Klinik und Poliklinik II, Lehrstuhl für Zelluläre Immuntherapie, Universitätsklinikum Würzburg, Würzburg, Germany

**Keywords:** Tumor-on-chip, Microphysiological systems, Immunotherapy, In vitro models, CAR-T cell therapy

## Abstract

The non-clinical assessment of CAR-T cells demands innovative models that are capable of predicting safety and efficacy in the clinical setting. Here, we present a novel solid tumor-on-chip model that allows CAR-T cell perfusion and integrates the vasculature and tumor lesions to recapitulate key events of CAR-T cell performance including extravasation, tumor infiltration and cytokine release. We assessed CAR-T cells targeting the ROR1 antigen against tumor aggregates that were derived from a breast cancer cell line and primary breast cancer organoids. The data show the temporal kinetic of ROR1 CAR-T cell migration and expansion, lytic activity and cytokine production over the course of 8 days, and reveal a correlation between anti-tumor efficacy and ROR1 antigen density on tumor cells. CAR-modified T cells extravasated faster, infiltrated tumor lesions stronger, persisted longer and in higher numbers than non-CAR modified T cells. Intriguingly, we detected cytokine release levels and kinetics typically observed in patients who developed cytokine release syndrome, and administered dasatinib as a pharmacologic OFF switch to control this inflammatory response. The data illustrate the ability of this tumor-on-chip platform to assess parameters associated withherapeutic outcome and the potential to aid in patient stratification and monitoring of CAR-T cell therapy.

## INTRODUCTION

Engineering T cells to express chimeric antigen receptors (CARs) has unveiled new opportunities in treating advanced-stage cancers, where T cells are used as a “living drug” acting directly on the tumor ^1^. Still, CAR-T cell therapy is limited to a small subset of cancer patients, particularly those with liquid tumors. In solid tumors, the efficacy of CAR-T cells is hindered by the immunosuppressive tumor microenvironment, high antigen heterogeneity and mechanisms involving tumor escape from immunosurveillance ^2^. Additionally, safety concerns such as cytokine release syndrome (CRS) are potentially life threatening and remain one of the major challenges in the clinical application of the therapy. CRS is characterized by the increase of cytokine levels in peripheral blood upon CAR-T cell infusion, which can lead to vascular damage in multiple tissues, organ dysfunctions, co-initiate immune effector cell-associated neurotoxicity syndrome (ICANS), and eventually patient death ^3^. The incidence rate of CRS ranges between 42-100%, and the severity can vary between patients ^4^. Multiple factors affecting the occurrence of CRS include tumor burden, CAR-T cell dose, and individual immune status, among others ^5^. This adverse reaction still mainly positions CAR-T cell therapy as a last-resort treatment option ^6^.

So far, nonclinical (mouse) models failed to predict these severe adverse effects, and other current models frequently fail to accurately predict clinical efficacy in solid tumors ^7, 8^. To investigate the mechanisms of adverse reactions and therefore enable the use of this therapy to its full potential, i.e., also in solid tumors, fit-for-purpose complex in vitro models are required. Such models should translate to human immunity, recapitulate tumor heterogeneity, its microenvironment and should therefore be able to better predict clinical safety and efficacy outcomes ^7^. Organ-on-Chip(OoC)-technologies already demonstrated the potential to fulfil those requirements, specifically by recapitulating tumor microenvironmental aspects^9^ and the integration of human immune components ^10, 11^.

Here, we demonstrate the application of a tumor-on-chip for the efficacy and safety assessment of ROR1-targeting CAR-T cells. We model the initial events upon CAR-T cells administration into the patients (CAR-T cells perfusion through the vasculature and extravasation towards the tumor) as well as the effects manifesting over the following week (CAR-T cell infiltration and migration as well as cytokine release kinetics). To demonstrate the opportunities our tumor-on-chip model provides, we further assessed its amenability to test combinational therapies by utilizing dasatinib as a CRS intervention strategy as well as its capability to include patient-specific effects by integrating patient-derived cancer organoids (PDOs).

## RESULTS

Concept and generation of the tumor-on-chip allowing CAR-T cells perfusion The tumor-on-chip is designed to mimic the solid tumor microenvironment and its interface with the vasculature, allowing the continuous perfusion of (CAR-)T cells. This allows the monitoring of (CAR-)T cell recruitment, infiltration into the tumor, and of tumor-(CAR-)T cell interactions and cytokine release kinetics over at least 8 days. Stacked microchannels separated by a porous membrane allow endothelial cell barrier formation and (CAR-)T cell perfusion through the top media channel (Fig. 1a, b) and placement of hydrogel-embedded tumor aggregates, fibroblast spheroids or patient-derived cancer organoids (PDOs) in a fixed position and within six individual cylindrical chambers (Fig. 1b).

**Figure 1:**
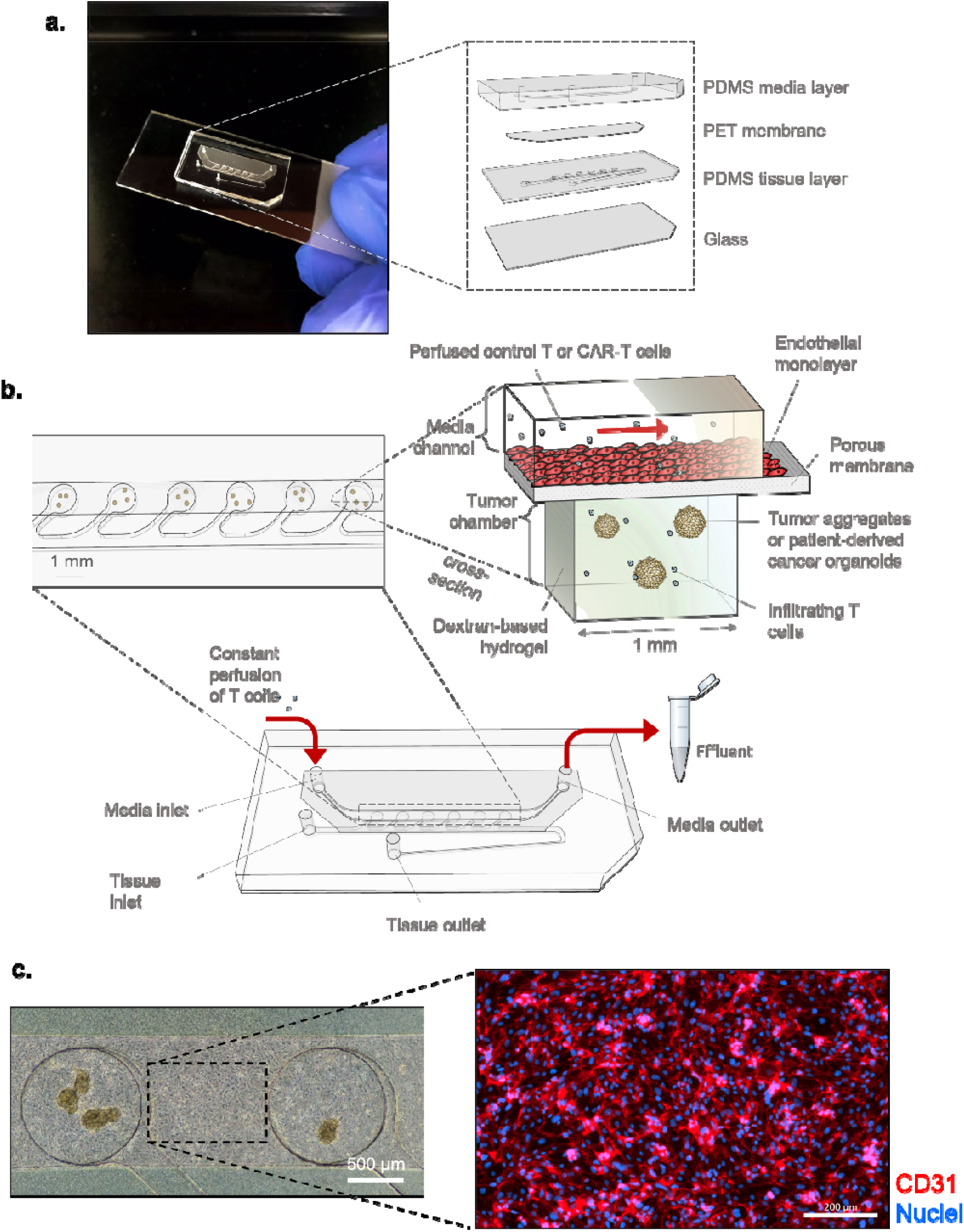
Concept and design of the tumor-on-chip. **a,** Tumor-on-chip with an exploded-view of the individual chip layers and materials. PDMS media layer contains the media channel and the fluidic inlets and outlets. PDMS tissue layer contains the tissue channel and chambers, which are separated from the media channel by a PET membrane and sealed at the bottom part with a microscope glass slide. **b,** Schematic representation of the tumor-on-chip model: cancer organoids/aggregates/fibroblast spheroids are embedded in a hydrogel and cultured in the chips’ tumor chamber, with six chambers in total per chip. The tumor chambers are located underneath a media channel, which is lined with an endothelial monolayer. Control T or CAR-T cells are perfused through the media channel and able to migrate into the tumor chamber. c, Brightfield image of the tumor-on-chip’s channel on day 0 before T cell perfusion. The media channel is covered by a tight monolayer of endothelial cells, visualized by CD31 immunofluorescence staining. Cell nuclei are shown in blue.

The typical experimental timeline for CAR-T cell treatment on the tumor-on-chip systems is illustrated in Fig. 2a. Following the injection of the target cells in the bottom chambers, either primary microvascular (allogeneic) or human induced pluripotent stem cell (hiPSC)-derived endothelial cells (ECs) sourced from the same (CAR-)T cell donor (isogenic) were seeded into the media channel and allowed to adhere overnight in a static condition before connecting the chips to constant media perfusion (20 µL/h). All tumor-on-chips included endothelial cells, if not indicated otherwise. The tumor-on-chips were perfused with either ROR1-targeting CAR-T cells or untransduced T cells from the same healthy donor as the control (referred to “control T cells” in the following). The perfusion was initiated once a tight endothelial barrier in the media channel was formed, characterized by CD31 expression (Fig. 1c). In all experiments, day 0 indicates the starting point of (CAR-)T cells perfusion. The perfusion process and parameters assured a 70-80 % T cell viability in both acellular chip and tumor-on-chip (Fig. 2b).

**Figure 2:**
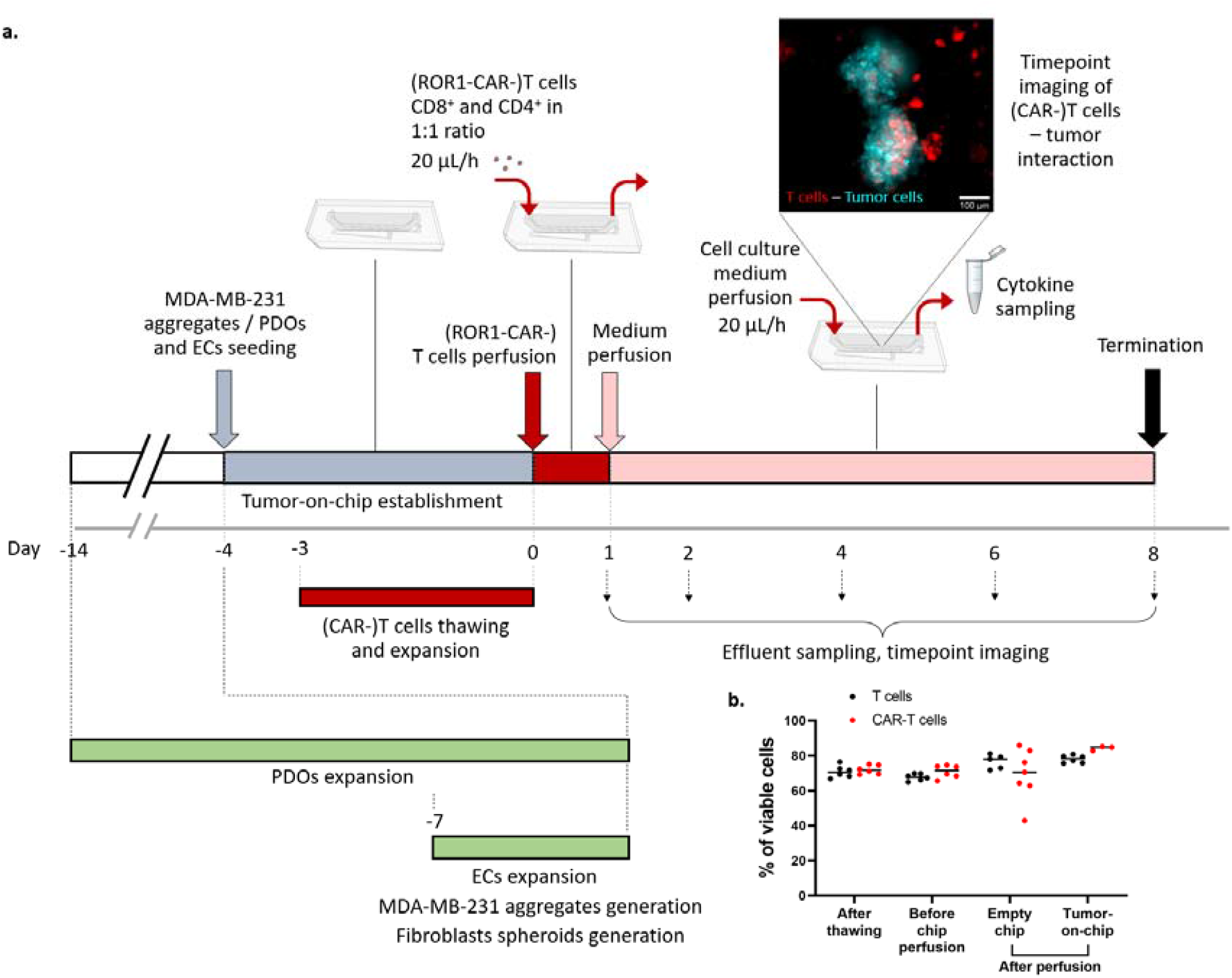
Timeline of the tumor-on-chip model generation and CAR-T cell treatment. **a,** Target cells—either MDA-MB-231, fibroblasts spheroids, or PDOs—, as well as endothelial cells were expanded and generated off-chip prior to chip seeding. Cells were seeded and cultured in the tumor-on-chip for a few days in parallel with the thawing and expansion of control T and CAR-T cells. At day 0, either control T or CAR-T cells were perfused through the chip for 20 h, followed by medium perfusion. Chips were imaged and effluents were collected every one or two days for up to day 8. **b,** Viability evaluation of the control T and CAR-T cells after thawing, before chip perfusion, and after perfusion either through empty chip or tumor-on-chip; n = 3-7.

### Modelling CAR-T cell recruitment, persistence, and tumor growth prevention on chip

To monitor antigen-specific antitumor activity of CAR-T cells, the recruitment of control T and CAR-T cells was measured in the presence of the microvascular endothelial cells (mvECs) barrier. On day 1, both control T and CAR-T cells infiltrated the GFP-expressing MDA-MB-231 tumor aggregates, with the CAR-T cells showing a significantly higher (2.5-fold) infiltration density (Fig. 3a, b). CAR-T cells also migrated towards the tumor aggregates faster than control T cells within the first one hour of perfusion (Suppl. Fig. 2a). Continuing the chip culture for up to 8 days, CAR-T cells persisted within the tumor aggregates and hampered their growth. On the other hand, the density of the infiltrating control T cells decreased over time and the tumor aggregates continued to grow (Fig. 3a, b, c). Control T cells were excluded to the outer area of most tumor aggregates on day 8 (Fig. 3a, Suppl. Fig. 2b). To quantify differences in the tumor mass, we measured GFP intensity of the MDA-MB-231 aggregates that showed an intensity drop in the CAR-T cell condition on day 1 but remained unchanged in the control T cell condition (Fig. 3d). When the (CAR-)T cells were perfused in lower concentration, i.e. 100,000 cells/mL, we observed less (CAR-)T cell migration into the tumor chamber (Fig. 3f). Nevertheless, the number of CAR-T cells that migrated into the tumor chamber remained significantly higher compared to the control T cells condition. Also in this lower concentration setting, no migration was observed when no tumor aggregate was present (Fig. 3e).

**Figure 3:**
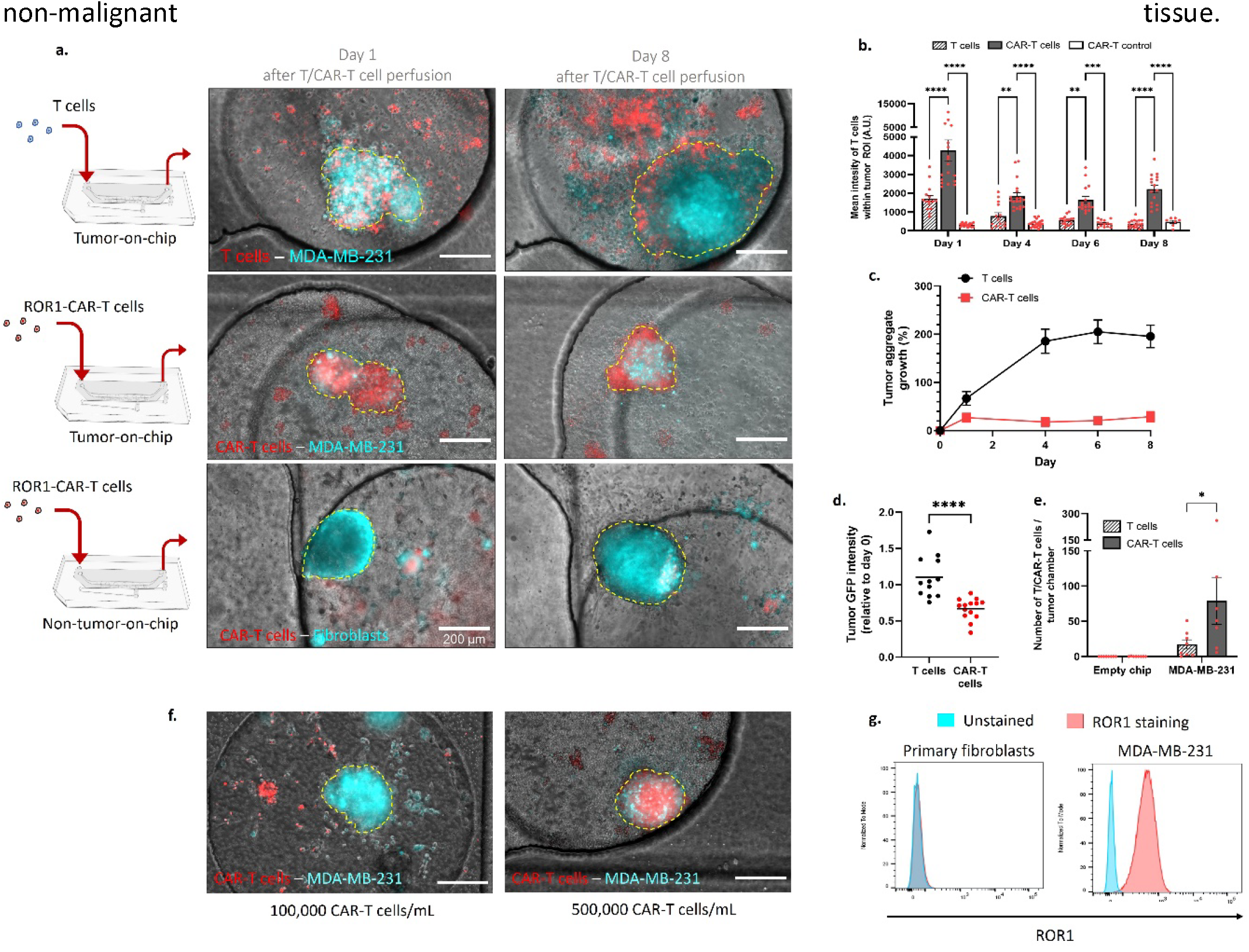
(CAR-)T cell infiltration and tumor aggregate growth upon (CAR-)T cell treatment. **a,** Representative images on day 1 and 8 of the MDA-MB-231 aggregates or fibroblast spheroids after tumor-on-chip perfusion with either control T or CAR-T cells, showing different degrees of T cell infiltration within the tumor chamber. MDA-MB-231 tumor cells express GFP and are pseudo-colored in blue, fibroblasts—representing non-malignant tissue—were labeled with CellTracker^TM^ CMFDA and pseudo-colored in cyan, whereas T and CAR-T cells were labeled in CellTracker^TM^ Deep Red. Yellow-dashed line marks the region of interest of each tumor aggregate/fibroblasts spheroid. MvECs were present in all chips. Scalebars: 200 µm. **b,** Quantification of mean intensity values of the control T and CAR-T cells within each tumor aggregate’s/fibroblast spheroid’s region of interest at different time points after (CAR-)T cell perfusion. “CAR-T control” indicates the condition where CAR-T cells were perfused through chips containing ROR1^−^ fibroblasts spheroids; n = 7-21 MDA-MB-231 tumor aggregates/fibroblast spheroids from 3-4 chips. Data are depicted as mean with ± SEM. **p<0.01, ***p<0.001, ****p<0.0001 as assessed by Bonferroni’s multiple comparisons test. **c,** MDA-MB-231 tumor aggregate growth post CAR-T cell treatment in comparison to (control) T cell condition as measured by quantifying the difference of each MDA-MB-231 aggregate area at different time points compared to their initial area before (CAR-)T cell perfusion; n = 12-16 aggregates from 4 chips. Depicted are mean ± SEM. **d,** Quantification of mean GFP intensity fold change expressed on each MDA-MB-231 tumor aggregate on day 1 after the perfusion of either (control) T or CAR-T cells. Each dot represents one tumor aggregate and black line indicates the mean value; n = 12-14 aggregates from 4 chips. ****p<0.0001; two-tailed unpaired t test. **e,** The bar graph shows the quantification of control T or CAR-T cells per tumor chamber on day 1 after the tumor-on-chip was perfused at a concentration of 100,000 (CAR-)T cells/mL. Empty chip was used as control; n = 7-8 chambers from 3 chips. Data are depicted as mean with ± SEM. *p<0.05; Bonferroni’s multiple comparisons test. Scalebar: 200 µm. **f,** Representative images on day 1 after perfusing the CAR-T cells (red) at either 100,000 or 500,000 cells/mL concentration through the tumor-on-chip containing MDA-MB-231 aggregates (green). The focal plane was set on the aggregates for all the acquisitions. Scalebar: 200 µm. **f,** Flow cytometry histograms showing ROR1 expression (red) in the MDA-MB-231 cells and fibroblasts compared with the unstained control (cyan).

### Probing off-target effects of CAR-T cells on chip

To assess off-tumor effects, CAR-T cells were perfused through chips containing ROR1^−^ fibroblast spheroids instead of ROR1^+^ MDA-MB-231 aggregates (Fig. 3a, g). Despite the migration into the bottom chamber, the fluorescence intensities of CAR-T cells within the fibroblast spheroids’ region remained low compared to the MDA-MB-231 aggregates over one week of culture period (Fig. 3a, b), demonstrating that ROR1-expression promoted CAR-T cells’ infiltration. Additionally, only low levels of IL-2, IL-6, IL-10, TNF-α, IFN-γ, and granzyme B were detected in this condition after 20 h of CAR-T cells’ perfusion (Suppl. Fig. 3a). This, therefore, demonstrated ROR1-specific CAR-T cell reactivity and the ability of the model to evaluate possible off-target effects by integrating cells from non-malignant tissue.

### Modelling CAR-T cell-mediated cytokine release

To model CRS on the tumor-on-chip and monitor clinically relevant inflammatory cytokine levels while simultaneously ensuring that it originates from the CAR-T cell – tumor interaction, we perfused either CAR-T or control T cells through acellular (empty), endothelial cell barrier (mvECs)-only, or complete tumor-on-chips, which included both mvECs and MDA-MB-231 tumor aggregates. The mvECs were ensured to express no ROR1 to exclude on-target off-tumor effects (Suppl. Fig. 1e). Compared to the T cell control, a higher level of all quantified cytokines, including IL-2, IL-6, IL-10, TNF-α, IFN-γ, and serine protease granzyme B, ranging from 6 to 1000-fold increase was detected on day 1 in the CAR-T cell condition when the tumor aggregates were present (Fig. 4a). The cytokine levels remained low in conditions without tumor aggregates (endothelial cells-only) and in acellular chips, showing that the cytokine secretion was induced by target antigen recognition (Fig. 4a).

**Figure 4:**
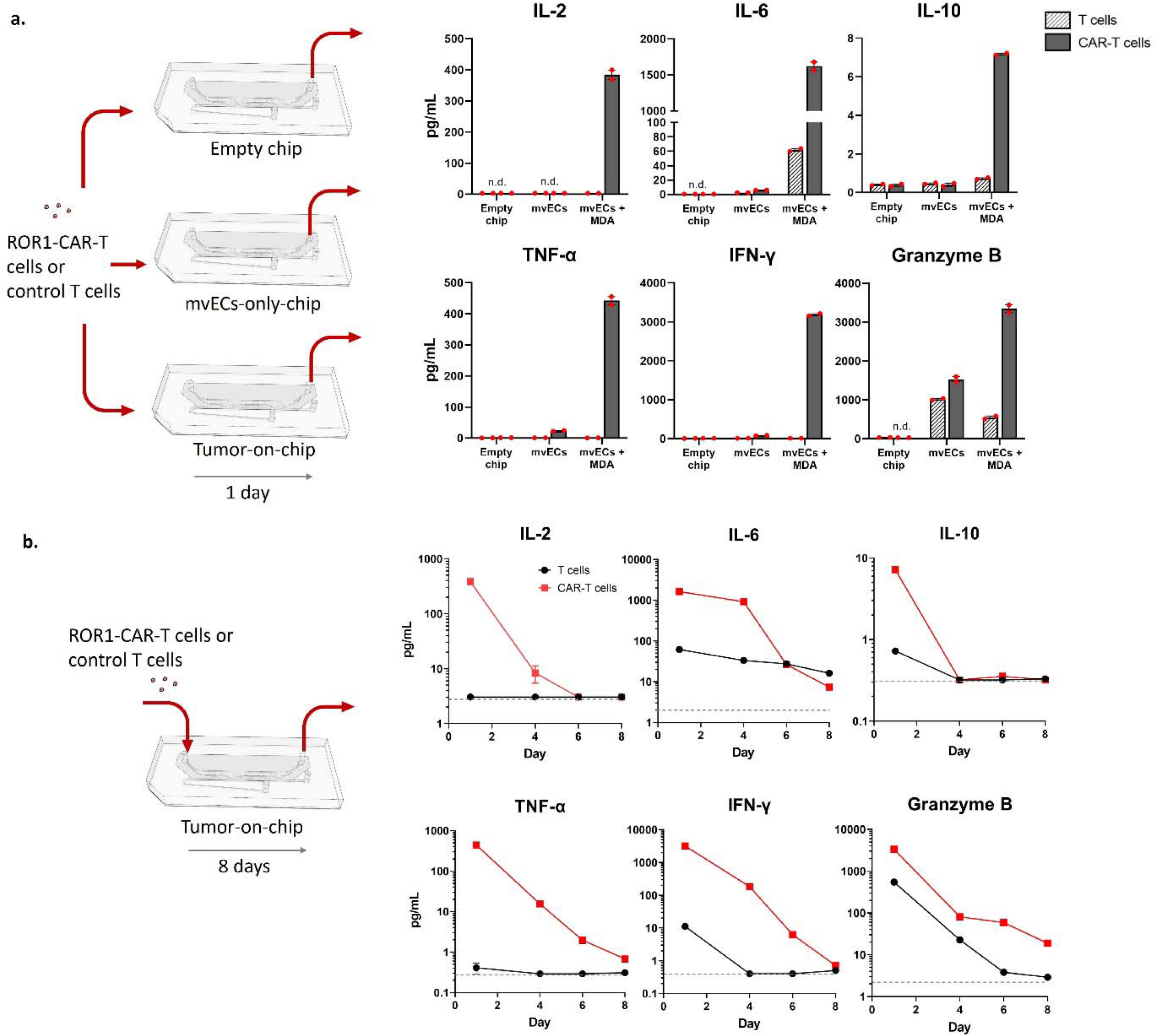
CAR-T cell-mediated cytokine release and its kinetics in the tumor-on-chip model. **a,** Quantification of the levels of cytokines IL-2, IL-6, IL-10, TNF-α, IFN-γ and granzyme B in the effluents of the chips after 20 h of (control) T or CAR-T cells perfusion through either acellular (empty) chips, mvECs-only, or complete tumor-on-chips (mvECs + MDA); n = 2 chips. Data are depicted as mean with ± SEM. Each red dot represents one chip. n.d.: not detected (below detection limit). **b,** Cytokine release kinetics of the abovementioned cytokines from day 1 to 8 after (control) T or CAR-T cells perfusion from day 0 to 1. Samples were collected every 2-3 days; n = 2 chips. Data are depicted as mean with ± SEM. The dashed line indicates the lower limit of detection.

The kinetics of cytokine release are depicted by collecting effluents over 8 days after (CAR-)T cell perfusion that lasted for 20 h. This longitudinal analysis of cytokine release showed the highest peak of cytokine concentration in the CAR-T cell condition on day 1 from all measured cytokines listed above, and consistently above the control T cells levels at almost all measurement time points (Fig. 4b). The drop in cytokine levels after day 1 might result from the ongoing lysis of MDA-MB-231 tumor aggregates by the CAR-T cells, as the tumors stopped growing in this condition as shown in the previous efficacy testruns (Fig. 3d). Using this setup, we were able to recapitulate clinically-relevant cytokine levels and kinetics mimicking patients’ condition suffering from high-grade CRS after receiving CAR-T cell therapy ^3^. Next, we further tested if the kinetics of cytokine secretion can be pharmacologically altered to model therapeutic intervention to manage the occurrence of CRS.

### Modelling CRS intervention on chip

CRS is highly challenging to predict so far ^7^. Once it occurs, using a pharmacological intervention that allows to temporarily “pause” CAR-T cell activity upon the onset of this immune-related adverse event could save patients from severe adverse reactions ^12^. Dasatinib is an FDA-approved tyrosine kinase inhibitor that inhibits phosphorylation of CD3ζ and ζ-chain of T cell receptor-associated protein kinase 70 (ZAP70) signalling in CAR constructs containing either CD28_CD3ζ or 4-1BB_CD3ζ activation modules in a reversible manner. We quantified the effects of introducing dasatinib to the CAR-T cell-treated tumor-on-chip either from day 1, from day 0 or only between days 0-1 to model CRS treatment, prevention, and delay, respectively (Fig. 5a).

**Figure 5:**
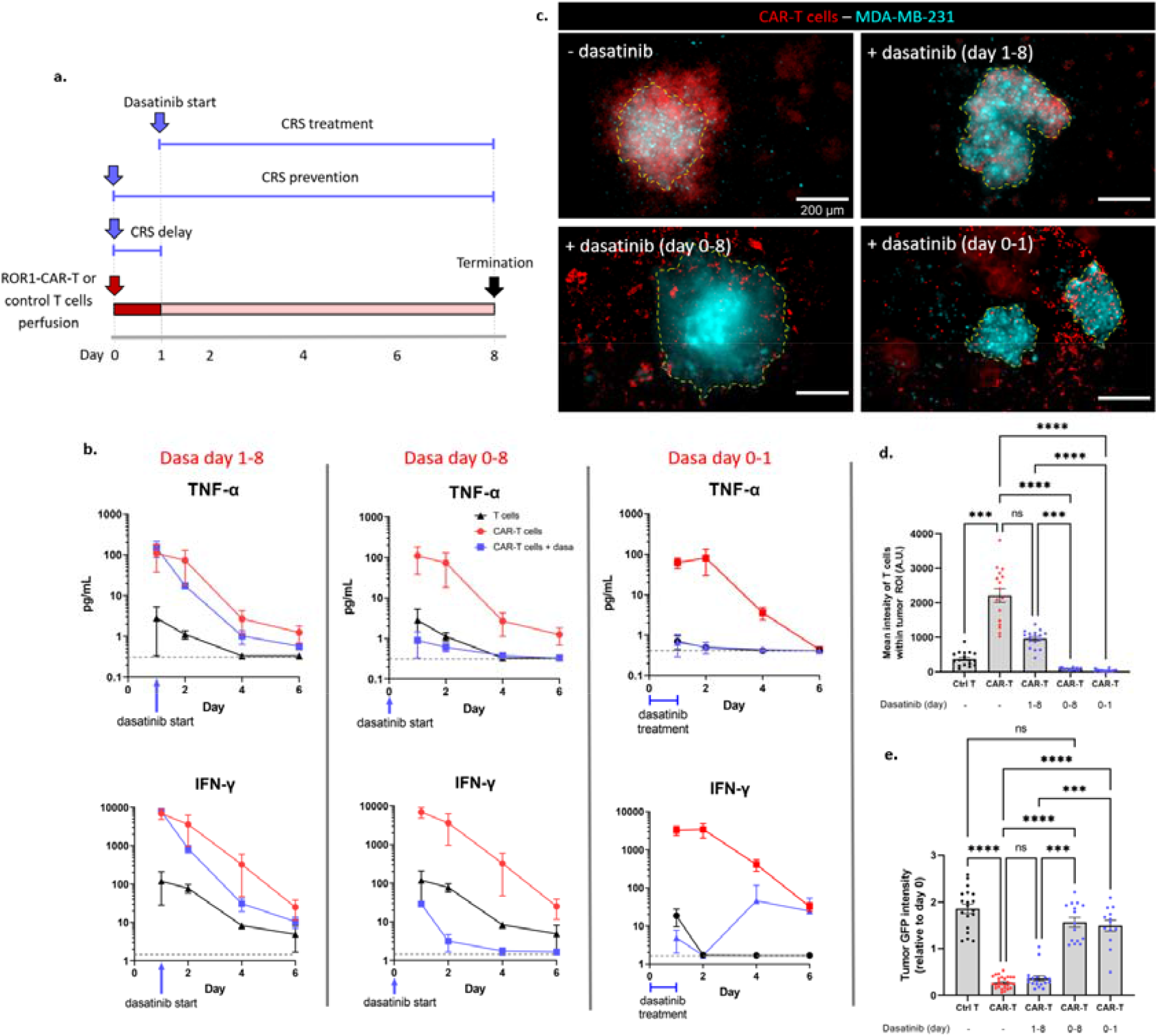
Dasatinib treatment to model CRS intervention strategy. **a,** Timeline of the dasatinib treatment with varying starting time point (blue arrow) and duration (blue bar) during the tumor-on-chip culture. Tumor-on-chips containing MDA-MB-231 aggregates were perfused on day 0-1 with either (control) T or CAR-T cells and subsequently with cell culture medium from day 1-8. 50 nM dasatinib was supplemented in the T / CAR-T cell suspension (day 0-1) and/or medium (day 1-8) depending on the experimental condition. **b,** TNF-α and IFN-γ release kinetics in control T, CAR-T cells and CAR-T cells + dasatinib condition in three different scenarios of intervention strategy: CRS treatment (left), prevention (middle) and delay (right). Data are shown as mean with ± SEM. Dashed line indicates the lower limit of detection. Data points from day 8 are excluded as the values are mostly below the detection limit. _n_ = 4 chips. **c,** Representative images on day 8 of the MDA-MB-231 aggregates (cyan) after tumor-on-chip perfusion with CAR-T cells (red), showing different degrees of T cell infiltration into the aggregate after different time point conditions of dasatinib treatment (day 1-8, 0-8, and 0-1) compared to untreated. Yellow-dashed line marks the region of one tumor aggregate. Scalebar: 200 µm. **d,** Quantification of mean intensity values of the control (Ctrl) T and CAR-T cells within each MDA-MB-231 tumor aggregate’s region of interest on day 8 after dasatinib treatment; _n_ = 15-18 MDA-MB-231 aggregates from 3-4 chips. Data are depicted as mean with ± SEM. ns: not significant, ***p<0.001, ****p<0.0001; Dunn’s multiple comparisons test. e, Quantification of mean GFP intensity fold change expressed on each MDA-MB-231 tumor aggregate on day 8 relative to mean GFP intensities of the aggregates on day 0. Each dot represents one MDA-MB-231 aggregate. Data are shown as mean with ± SEM; n = 13-28 aggregates from 3-4 chips. ns: not significant, ***p<0.001, ****p<0.0001; Dunn’s multiple comparisons test.

Starting dasatinib application on day 1—at the peak of cytokine levels seen in previous experiments (Fig. 4)—resulted in a fast drop of TNF-α and IFN-γ levels relative to the untreated CAR-T cells condition, thereby mimicking acute management of CRS. This treatment scheme also enabled the CAR-T cells to infiltrate the tumor within the first 24 hours but then stopped the subsequent excessive infiltration, as indicated by lower T cell intensity (∼2-fold) compared to the untreated CAR-T cell condition (Fig. 5c, d). Furthermore, the tumor GFP signal intensity on day 8 dropped to a similar level as the CAR-T cell condition without dasatinib (Fig. 5e), thereby maintaining their base antitumor cytotoxicity.

In contrast to the day 1-8 treatment, the day 0-8 treatment approach prevented the cytokine release, as shown in the low TNF-α and IFN-γ levels compared to the untreated CAR-T cells and control T cells conditions (Fig. 5b). However, this consequently also prevented the CAR-T cells to infiltrate the tumor and the signal intensity from the GFP-expressing MDA-MB-231 tumor aggregates was minimally changed, with no significant differences compared to the control T cells condition (Fig. 5d, e). This, therefore, defeats the purpose of CAR-T cell treatment. Day 0-1 treatment delayed IFN-γ but not TNF-α peak, showing a transient effect of dasatinib (Fig. 5b). However, this approach also prevented the CAR-T cells to infiltrate and lyse the tumor aggregates as seen on day 8 (Fig. 5c, d, e).

By investigating different intervention time points and durations in the CAR-T cell-treated tumor-on-chips, we hereby demonstrated the feasibility to determine a suitable CRS intervention strategy to achieve maximum safety without diminishing the efficacy of CAR-T cell therapy.

### Modelling patient-specific effects using patient-derived cancer organoids

ROR1 expression levels are tumor- and patient-specific. To validate the therapeutic efficiency within this heterogeneity, we used patient-derived tumor organoids (PDOs) expressing high and low ROR1 levels derived from two patients (Fig. 6a, Suppl. Tab. 1), and compared with MDA-MB-231 aggregates as a positive control condition for CRS emulation. To capture these tumor- and patient-specific effects exerted from the PDOs, possible alloreaction of T cells towards the endothelial cells—e.g. when primary allogeneic mvECs were employed—were excluded by integrating hiPSC-derived ECs sourced from the same donor as the (CAR-)T cells.

**Figure 6:**
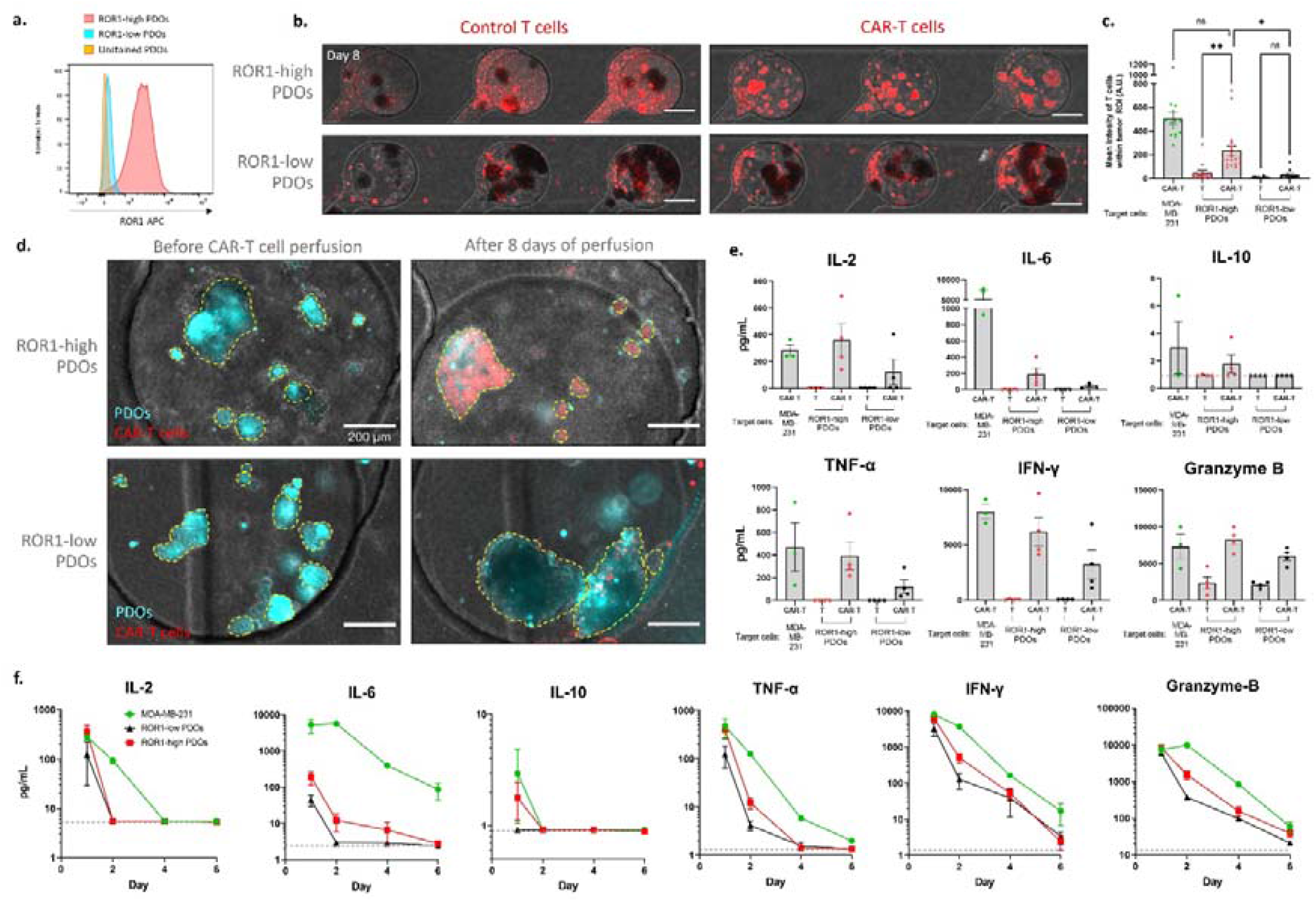
CAR-T cells infiltration and cytokine secretion towards patient-derived cancer organoids. **a,** Flow cytometry histogram plots showing ROR1 expression in the high-ROR1-expressing PDOs (red), low-ROR1-expressing PDOs (cyan) compared with the unstained control (orange). **b,** Representative images on day 8 of the ROR1-low and -high expressing PDOs after perfused with either (control) T or CAR-T cells. Fluorescently-labeled T cells are pseudo-colored in red. The focal plane was set on the PDOs for all the acquisitions. Scalebar: 500 µm. **c,** Quantification of mean intensity values of the control T (depicted as “T” in the graph) and CAR-T cells within the region of interest of each PDO on day 8. MDA-MB-231 aggregates were used as positive control; n = 14-20 PDOs from 3-4 chips. Data are depicted as mean with ± SEM. ns: not significant, *p<0.05, **p<0.01; Dunn’s multiple comparisons test. **d,** Representative images of the ROR1-high and ROR1-low expressing PDOs (cyan) before CAR-T cell (red) perfusion and on day 8 of the experiment. Yellow-dashed line marks the region of interest of each PDO. Scalebar: 200 µm. **e,** Quantification of the levels of cytokines IL-2, IL-6, IL-10, TNF-α, IFN-γ and granzyme B in the effluents of the chips after 20 h of (control) T or CAR-T cells perfusion through the tumor-on-chip containing either MDA-MB-231 aggregates (positive control for CAR-T cell treatment), ROR1-high PDOs or ROR1-low PDOs; n = 3-4 chips. Data are depicted as mean with ± SEM. Each red dot represents one chip. **f,** Cytokine release kinetics of the abovementioned cytokines from day 1 to 6 after CAR-T cells perfusion from day 0 to 1 through the tumor-on-chip containing either MDA-MB-231 aggregates (positive control for CAR-T cell treatment), ROR1-high PDOs or ROR1-low PDOs; n = 4 chips. Data are depicted as mean with ± SEM. Dashed line indicates the lower limit of detection. Data points from day 8 are excluded as the values are mostly below the detection limit.

In the semi-autologous setup of the tumor-on-chip, a dose-response depending on the ROR1 expression level was observed for the CAR-T cell infiltration into the tumor (Fig. 6a-d) as well as the cytokine secretion level (Fig. 6e, Suppl. Fig. 5) and long-term cytokine kinetics (Fig. 6f). In correlation with their respective target expression level, CAR-T cells were infiltrating the ROR1-high PDOs in significantly higher numbers (9-fold) compared to ROR1-low PDOs. The highest CAR-T infiltration rate was observed into MDA-MB-231 aggregates: 2-fold higher compared to the ROR1-high PDOs condition (Fig. 6c). Additionally, ROR1-low PDOs merged over time and continued to grow despite CAR-T cell treatment, whereas the growth of ROR1-high PDOs was hampered (Fig. 6d).

It is worth noting that despite the absence of CAR, control T cells were still recruited into the tumor chamber, which might indicate a baseline allogeneic response towards the PDOs. However, control T cells mostly remained excluded from the PDOs after the culture period (Fig. 6b) and resulted in much lower cytokine levels on day 1 than the respective CAR-T cell conditions (Fig. 6e).

## DISCUSSION

This tumor-on-chip model demonstrates for the first time the possibility to assess both the efficacy and safety aspect of a CAR-T cell therapy for solid tumors in vitro. Our model recapitulates multiple phases of the therapy, including CAR-T cells’ recruitment from the blood vessel into the tumor site upon therapy administration followed by multiple-days T cells-tumor interaction within the tumor microenvironment; here demonstrated by CAR-T cells targeting ROR1. ROR1 is expressed in many epithelial tumors and a subset of B cell tumors ^13, 14^. In non-malignant adult tissues, ROR1 expression is absent except in B cells precursors that undergo a specific stage of maturation in the bone marrow^13, 15^. As MDA-MB-231 cells overexpress ROR1 antigen, they were selected as the target cells in our experiments. Human primary fibroblasts were used to represent non-malignant tissue that lacks ROR1 expression (Fig. 3g).

By perfusing the (CAR-)T cells through the vasculature-like channel, we were able to assess their recruitment into the tumor site and observed that CAR-T cells extravasated faster, infiltrated the MDA-MB-231 aggregates in higher density, and persisted longer than unmodified T cells. We observed the exclusion of unmodified control T cells from the tumor aggregates after one week of culture (Suppl. Fig. 2b)—an observed clinical phenomenon to predict the subset of cancer patients who are responsive to cancer immunotherapy ^16, 17^. Further, we found that the relative ROR1 expression on the PDOs positively correlates with the persistence of CAR-T cells within the tumor bulk, which is aligned with recently published data, where antitumor-killing of breast tumor-derived PDOs within the first 24 h positively correlates with the ROR1 expression level ^18^. The integration of PDOs may therefore recapitulate target-antigen heterogeneity and patient-specific response, which has recently been reported in multiple studies that mainly focus on the efficacy aspect ^18–22^.

To capture patient-specific effects exerted from the PDOs, we refined the model by integrating isogenic hiPSC-ECs. As demonstrated in our cytokine secretion data, higher fold change values of TNF-α, IFN-γ and granzyme B level were detected when T cells were co-cultured with mvECs from multiple donors compared to the co-culture condition with the isogenic hiPSC-ECs. The fold change of cytokine level in isogenic setting remained similar to the condition without endothelial cells (Suppl. Fig. 4). HiPSC-ECs were ensured to express CD31 prior to integration into the chip (Suppl. Fig. 8b). Despite enabling human leukocyte antigen (HLA)-matched autologous systems, hiPSCs-ECs expressed a certain level of ROR1 (Suppl. Fig. 8a), which induced IFN-γ and granzyme B secretion in our plate assay (Suppl. Fig. 8c). Indeed, ROR1 is not only a tumor antigen but also a known marker for stem cell derivatives ^23^. Despite the resulting background activation, patient-specific effects from the relative ROR1 expression levels of the PDOs could be detected in the chips. For different types of CARs, this aspect should not apply and future advances in EC maturation should allow to diminish this effect also for ROR1-CAR-T cells.

In terms of safety, the elucidation of the toxicity mechanisms of CRS and prediction of patient response urgently requires novel models ^7, 9, 24^. Current in vitro models are unable to follow the kinetics of cytokine secretion for safety evaluation ^7^. To address this aspect in our chip system, we enabled linear perfusion of (CAR-)T cells followed by cell culture medium with a constant flow rate (20 µL/h) through the chips for over one week. Here, we first tested three different perfused CAR-T cell concentrations (100,000, 500,000, and 1,000,000 T cells/mL) while keeping the number of MDA-MB-231 tumor aggregates relatively constant. We selected the values based on the range of peripheral T cell count in patients who had cytokine release syndrome after receiving CD19-CAR-T cell therapy, and proceeded with 500,000 T cells/mL for subsequent experiments as this produced clinically-relevant and detectable cytokine levels (Suppl. Fig. 3b) ^25, 26^. Moreover, a concentration higher than 500,000 T cells/mL often led to T cell clumping in the media channel, which consequently disrupted the fluidic flow.

Upon CAR-T cell administration, we consistently detected peaks of cytokine level on day 1. The level typically decreased afterwards, and it remains to be investigated whether this was caused by an activation-induced cell death of T cells following eradication of the tumor cells or T cell exhaustion ^3^. Additionally, replacing the non-recruited CAR-T cells with cell culture medium in the perfusate from day 1 may limit the cytokine secretion only from the recruited CAR-T cells in subsequent days. Nevertheless, the presence of granzyme B was still detected on day 8 of culture in both CAR-T and control T cell conditions, indicating an ongoing T cell-tumor interaction. Early secretion—up to day 3 after CAR-T cell administration—of IFN-γ was shown to predict patients who will develop severe CRS^27^. Circulating IFN-γ level was associated with CRS in 12 out of 13 trials ^28^. The concentration of IFN-γ typically found in CRS patients (>10 pg/mL) is indeed much lower than the one detected from chips experiments (>3000 pg/mL) ^26, 28^. However, a 100-fold increase in IFN-γ serum concentration was detected in patients suffering from severe CRS ^3^, which could also be observed in our model even after 4 days of chip culture.

Interestingly, we noticed differences in the cytokine values depending on the CAR-T cell donor used for the perfusion, although the positive fold change values typically decreased to below 20 on day 6 for both CAR-T cell donors tested (Suppl. Fig. 7). This also demonstrates the model’s applicability for safety evaluation of different CAR-T cell products and potential use for patient stratification. Further, our data show a similar pattern of onset and kinetics of IL-2, IL-10, and TNF-α secretion as observed also in patients suffering from high-grade CRS, although on a shorter time scale ^3, 29^ Further tests and extensive benchmarking with clinical data (“real-world data sets”) may elucidate whether the absolute and fold change values from the chips could be graded as “toxic” or “non-toxic”.

Another cytokine, namely IL-6, has been identified as a crucial mediator of CRS ^30^. High IL-6 serum level are frequently detected in patients who suffer from severe CRS and blocking of IL-6 signalling (e.g. with tocilizumab) together with systemic immunosuppression via corticosteroids are the current standard of care in CRS management ^3, 31–37^. Although we saw an increase in IL-6 production in the CAR-T cell condition almost at all time points and for different donors, it is important to note that our model did not include endogenous innate immune cells such as macrophages and monocytes, which are the main source of IL-6 that eventually induces systemic inflammation ^38^. Their future incorporation will therefore be crucial in further understanding their role in the amplification and perpetuation of the inflammatory cycle that would lead to CRS. For this reason also, tocilizumab is arguably not the best option to test CRS intervention in our model, and interfering directly with CAR-T cells’ function, e.g. with dasatinib, provides a better alternative instead. Besides, hyperactivation and hyperproliferation of the CAR-T cells are seen as the root factor of the subsequently developing CRS ^4^. Applying dasatinib temporarily inactivates the CAR-T cells without negatively affecting their viability ^39^, thus may be applied to manage acute toxicity. In preliminary experiments using cell culture well plates, we initially confirmed that dasatinib (50 nM) did not negatively affect CAR-T cell viability and successfully inhibited the upregulation of early (CD69) and late (CD25) activation markers after 24 h interaction with MDA-MB-231 cells (Suppl. Fig. 6). By applying dasatinib at different time points and in varying duration, we demonstrated that an optimal strategy to control cytokine release while keeping the CAR-T cells efficacious throughout one week period can be identified.

Owing to the chip’s modularity, integrating additional components into the tumor microenvironment (e.g. ECM components, varying gel stiffnesses, and other biophysical forces) and/or applying different cell sourcing strategies (allogeneic, semi-autologous, fully autologous) may allow the study of their individual impact on CAR-T cell efficacy and safety. In a clinical setting, sourcing the cells directly from cancer patients receiving CAR-T cell therapy and monitor the development in the tumor-on-chip model may help clinical decision-making. As a part of the project imSAVAR (Immune Safety Avatar: nonclinical mimicking of the immune system effects of immunomodulatory therapies; www.imsavar.eu), one of our aims is to develop novel models that reflect mechanisms (i.e. key events and key event relationships within a specific immune-related adverse outcome pathway, irAOP) and conditions (e.g. the tumor microenvironment) for improved nonclinical safety assessment. Despite its potential and our comprehensive characterization, it will require an extensive benchmarking with clinical data to fully confirm that such complex tumor-on-chip models possess a higher predictive value for clinical safety and efficacy compared to animal models. Nevertheless, the development of complex, human-relevant tumor models offers new opportunities both for mechanistic research of CAR-T cell therapy and clinical translation of novel CAR-T cell products.

## MATERIALS AND METHODS

### Tumor chip concept, design and fabrication

The tumor-on-chip consists of two stacked microfluidic channels separated by an isoporous, semipermeable polyethylene terephthalate (PET) membrane (5 µm poresize: r_P_1=151µm; ρ 1=161×110^5^ pores per cm^2^; TRAKETCH® PET 5.0 p S2101×1300, SABEU GmbH & Co. KG, Northeim, Germany), which was functionalized by a plasma-enhanced, chemical vapor deposition (PECVD) process as previously described ^40^. The tumor chambers (1 mm in diameter, 0.3 mm in height, six chambers per chip) branch off a main injection channel (0.2 mm in height) at a 45° angle and a high-resistance channel towards the outlet port, which forces the aggregates/organoids to sequentially enter the tumor chambers during the injection process, utilizing an injection mechanism previously successfully established in other organ-on-chip model ^11^. This concept and design enable the integration of any aggregate/organoid below 0.2 mm in size. The tissue channel is designed to have an outlet channel that possesses a fluidic resistance of 1.2 x 10^12^ Pa.s/m^3^, which is at least twice higher than the fluidic resistance of the membrane and the media channel combined. This ensures the placement of the cells into the tumor chambers during the injection instead of flowing out towards the outlet. The media channel is situated right above the tumor chambers and is separated by a porous PET membrane that serves multiple functions: during the cell injection step, it enables the entrapment of the tumor aggregates/PDOs and provides a growth surface for the endothelial barrier formation, whereas during the culture period, its 5 µm pore size allows T cell trafficking and passive diffusion of diluted species ^11^.

The tumor tissue and media layers consisting 200-300 µm high channels were micro-structured via replica molding of polydimethylsiloxane (PDMS; Sylgard 184, Dow Corning, USA) on two differently patterned master wafers fabricated via photolithographic processes ^41^. The SU-8 structures on the tumor tissue layer wafer have two heights: 300 µm for the round tumor chambers and 200 µm for the rest of the channels. For the replica molding, PDMS was homogeneously mixed in a 10:1 (elastomer base:curing agent) mass ratio and then degassed in a desiccator to remove air bubbles. Afterwards, two different replica molding approaches were conducted as previously described ^11^. Standard molding approach was used by pouring the PDMS prepolymer solution onto the silicon wafer master mold to obtain 3 mm thick PDMS pieces with channel structures for the media layer. PDMS was cured at 60 °C for 4 h. After curing, the PDMS were cut to the size of the chip and ports were pierced using a biopsy punch (Disposable Biopsy Punch, 0.75 mm diameter; 504 529; World Precision Instruments, Friedberg, Germany) to access the chips (for both cell injection and medium perfusion). The exclusion molding approach was used to fabricate the bottom tumor layer: here, PDMS prepolymer solution was poured onto the silicon wafer master mold, which was then clamped against a 5 mm-thick PMMA disk to produce a 0.3 mm thin layer with through hole channel structures. PDMS was cured at 60 °C for 2 h. Once the PDMS parts were cured, they were cleaned using isopropanol followed by deionized water and blow-dried with nitrogen pistol. The microfluidic chips were assembled in three consecutive bonding steps: (i) tumor layer to glass, (ii) media layer to the membrane, (ii), and (iii) the tumor layer to media layer. In all steps, bonding was achieved by oxygen plasma activation (75 W, 0.2 cm^3^m−1 O ; Diener Zepto, Diener electronic GmbH + Co. KG, Ebhausen, Germany) for 24 s. Bonded parts were baked at 60 °C for at least 30 min after each bonding step, and overnight after the entire chip was assembled. All chips were O_2_-plasma treated (751W, 0.21cm^3^m−1 O) for 51min to sterilize and hydrophilize the PDMS surface before cell injection.

### Tumor and fibroblast aggregates generation

MDA-MB-231 cells were purchased from ATCC and were lentivirally transduced to express GFP-firefly luciferase fusion protein. The cells were cultured in RPMI-1640 (11835030, Thermo Fisher Scientific) supplemented with 10% (v/v) FCS (SH30066.03, HyClone™ FetalClone™ II Serum; Cytiva), 1X GlutaMAX (35050061; Thermo Fisher Scientific), and 100 U/mL penicillin-streptomycin (15140122; Thermo Fisher Scientific) in a humidified incubator at 37 °C, 5% CO_2_ and 95% rH. Aggregates were formed using agarose microwells-based approach ^42^. Briefly, 2.5 g of Hydrosil silicone components (1:1 wt) (101301; SILADENT) was added into each well of 6-well microwell culture plate (AggreWell^TM^400; STEMCELL Technologies) followed by centrifugation at 55 x g for 1 min and curing at 60 °C for 1 h. The cured molds were then carefully removed, cut into circular segments (d = 20 mm), and glued to a polymethyl methacrylate (PMMA) holder that has circular cutouts that match the surface of a 24-well plate (d = 15.5 mm, h = 2 mm). These cured molds were used as a reusable master mold for the agarose gel.

Prior to aggregate formation, the master molds were disinfected with 70% (v/v) ethanol and rinsed three times with PBS without Ca and Mg (indicated as PBS^−^ in the following) (P04-36500; PAN-Biotech). Afterwards, 3% (w/v) agarose solution in DMEM (41965039; Gibco) was liquified by preheating in a microwave and 650 µL was added into each master mold and left for 10 min at room temperature (RT) to allow agarose gelation. Afterwards, the agarose molds were transferred into the wells of 24-well plate with the patterned structures facing upwards. The wells were filled with 1 mL of PBS^−^ followed by centrifugation at 1300 x g for 3 min to remove trapped air bubble below the agarose mold. PBS^−^ was removed before adding the cells.

To generate the MDA-MB-231 aggregates, the cells were detached by incubation for 3 min with TrypLE^TM^ Express Enzyme (12604013; Gibco) at 37 °C, pelleted by centrifugation at 200 x g for 5 min and resuspended in RPMI-1640-based cell culture medium described above. The cell suspension was then mixed with basement membrane extract (BME) (Cultrex Reduced Growth Factor Basement Membrane Extract, Type 2 Select, 3533-005-02; Bio-techne) in 1:1 (v/v) ratio on ice and pipetted into each agarose well with 200 µL mixture solution containing 100,000 cells. The 24-well plate was immediately centrifuged at 4 °C at 900 x g for 10 min, followed incubation for 30 min at 37 °C, 5% CO_2_ and 95% rH for BME gelation. Afterwards, each well was supplemented with 1 mL of cell culture medium for 3 days of culture until chip injection. The aggregation method resulted in aggregates with the size of ∼120 µm after 3 days of culture (Suppl. Fig. 1a, b).

Primary human fibroblasts were cultured in DMEM (41965039; Gibco) supplemented with 10% (v/v) FCS and 100 U/mL penicillin-streptomycin. To generate the fibroblast aggregates, a similar approach as described above was followed: fibroblasts were detached using TrypLE^TM^ Express Enzyme for 3 min at 37 °C, pelleted by centrifugation at 200 x g for 5 min and resuspended in cell culture medium. 100,000 cells in 1 mL medium were pipetted into each agarose well. The 24-well plate was then centrifuged at RT at 900 x g for 10 min, and incubated at 37 °C, 5% CO_2_ and 95% rH for 3 days until chip injection.

### Patient-derived cancer organoids generation and culture

Patient-derived organoids (PDOs) were established from specimens obtained from advanced breast cancer patients treated at the Department of Women’s Health in Tübingen. All patients gave informed consent and the study was approved by the Ethical Committee of the Eberhard Karls University Tübingen (No. 150/2018BO2 and 662/2022BO2).

Pleural effusion (PE) samples from two patients were collected by thoracentesis and processed as followed. PE samples were centrifuged at 500 x g for 10 min. Cell pellets were pooled and when necessary, red blood cells were lysed with 10 mL of RBC lysis buffer (155 mM NH_4_Cl, 10 mM KHCO_3_, 100 µM Na2EDTA in H_2_O, pH 7.4) on ice for 5 min. Cells were diluted in DPBS (Dulbecco1s Phosphate Buffered Saline, P04-36500; Pan Biotech GmbH) and centrifuged at 500 x g for 10 min. The final cell pellet was resuspended in AdvDMEM/F12+++ (Advanced DMEM/F-12 (12534010), 1% Pen/Strep (15140122), 1x GlutaMAX (35050-038), 10 mM HEPES (15630-056); all from Thermo Fisher). For organoid culture setup, the desired amount of cell suspension was mixed with Basement Membrane Extract (BME; Cultrex Reduced Growth Factor Basement Membrane Extract, Type 2 Select, Bio-techne, 3533-005-02) at a ratio of 30% cell suspension to 70% BME. 20 µL droplets were plated into wells of a 48-well plate and placed upside-down in an incubator (37°C, 5% CO_2_) to solidify for 30 min. Afterwards, 280 µL of culture medium (AdvDMEM/F12+++ supplemented with 10% (v/v) conditioned medium from L-WRN cells (ATCC-CRL-3276) ^43^, 5 nM Heregulinß-1 (100-03; PeproTech, NJ, USA), 5 ng/mL fibroblast growth factor 7 (FGF7) (100-19; PeproTech), 20 ng/mL fibroblast growth factor 10 (FGF10) (100-26; PeproTech), 5 ng/mL epidermal growth factor (EGF) (AF-100-15; PeproTech), 500 nM A83-01 (2939; Tocris, Wiesbaden, Germany), 5 µM Y27632 (72034; STEMCELL Technologies), 500 nM SB-202190 (S7067; Sigma-Aldrich), 2% B27^TM^ Supplement (17504-044; Thermo Fisher Scientific), 1.25 mM N-acetyl-cysteine (A9165; Sigma-Aldrich), 5 mM nicotinamide (N0636; Sigma-Aldrich)) was added to each well. The medium was changed every 3–4 days.

The PDOs were passaged based on the confluency of the culture, ranging from 5-20 days. PDOs were recovered from the wells by resuspending the BME-droplets in ice-cold DPBS containing 5 µM Y-27632 (DPBS/Y-27632). This organoid suspension was incubated with 1 mL of TrypLE™ Express Enzyme (1X; Thermo Fisher, 12604013) at 37°C in a water bath for 5 min, followed by mechanical dissociation via 5 times pulling and dispensing the solution using a syringe with 27G needle (302200; Becton Dickinson). The suspension was then centrifuged at 500 x g for 10 min and the supernatant removed. For further culture, the desired amount of cell pellet was resuspended in AdvDMEM/F12+++ and mixed with BME at a ratio of 30% cell suspension to 70% BME and cultured as described above.

Prior to chip injection, the PDOs were expanded in a droplet of basement membrane extract hydrogel until they reached ∼120 µm in diameter, typically after ∼10 days. To anticipate patient-specific differences in growth speed, the PDOs culture was monitored daily to better estimate the appropriate loading day. To retrieve the PDOs, gel droplets were first washed with PBS^−^ before introducing 500 µL of 2 mg/mL cold dispase II solution followed by pipetting up and down to break the gel droplet. The organoid cell suspension was then incubated for 10 min at 37 °C, 5% CO_2_ and 95% rH. Afterwards, each well was resuspended with 500 µL of AdvDMEM/F-12+++ and the retrieved PDOs were centrifuged at 500 x g for 5 min. Each droplet of PDOs on day 10 of culture was used to load 3 chips. After pelleting the PDOs, they were incubated in 10 µM of CellTracker^TM^ CMFDA solution (dissolved in phenol red-free DMEM/F-12 medium) for 20 min at 37°C, 5% CO_2_ and 95% rH. The organoids were then washed twice by adding phenol red-free DMEM/F-12 medium up to 10 mL and centrifuged at 500 x g for 5 min for each washing cycle. The organoid pellet was then resuspended in dextran-based hydrogel as will be described in the next section.

### Microvascular endothelial cells isolation and culture

Human microvascular endothelial cells (mvECs) were isolated from resected skin tissue from plastic surgeries received from Dr. Ulrich E. Ziegler (Klinik Charlottenhaus, Stuttgart, Germany) as described previously ^11^, and approved by the local medical ethics committee: Patients gave an informed consent according to the permission of the “Landesärztekammer Baden-Württemberg” (IRB#: F-2020-166; for normal skin from elective surgeries). Subcutaneous fat, visible blood vessels, and connective tissue were removed, and the remaining skin tissues were cut into strips of ∼4 cm length and ∼1 mm width, followed by incubation in 10 mL of 2 U/mL dispase solution (D4693; Merck KGaA) at 4 °C overnight. Next, the epidermis layer was peeled off and discarded, and the remaining dermis layer was washed twice in PBS^−^. The mvECs were isolated from the dermis by incubating the dermis strips in 0.05% trypsin/EDTA (59417C; Sigma) for 40 min at 37 °C, followed by stopping the trypsinization with cell culture medium supplemented with 10% FCS and transferring the strips into a petri dish containing pre-warmed PBS^−^. Next, the dissociated cells were scraped out with a scalpel and the cell suspension was strained through a 70 µm strainer (542070 ; Greiner Bio-One) and collected. The cell suspension was then centrifuged at 209 x g for 5 min. Lastly, the cell pellet was resuspended in 10 mL of Endothelial Cell Growth Medium (ECGM, C-22010; PromoCell GmbH) supplemented with 10 mg/mL Gentamicin (15710049; Thermo Fisher Scientific). The cells were seeded into two T25 cell culture flasks (690175; Greiner Bio-One) and incubated at 37 °C, 5% CO_2_ and 95% rH. The cells were further expanded in T75 (658175; Greiner Bio-One) and used in passage 2 or 3 for this study.

### Human induced pluripotent stem cells generation from PBMCs

HiPSCs were generated from two female donor PBMCs using episomal vectors without TP53 shRNA^44^ according to standard protocols in the Leiden University Medical Center (LUMC) iPSC core facility and following informed consent at the University Hospital Clinic in Wurzburg, Germany. HiPSCs were routinely cultured on Vitronectin XF in TeSR-E8 (all from Stem Cell Technologies) according to the manufacturer’s protocol. Standard characterization of hiPSCs was performed as described previously^45–47^. Pluripotency of the hiPSC clones was confirmed by flow cytometry analysis for OCT3/4, SSEA-4, NANOG and spontaneous differentiation towards three germ lineages. G-banding analysis was conducted at the Laboratorium voor Diagnostische Genoomanalyse (LGDA), LUMC according to standard procedures. A total of 20 metaphases were analyzed for each line. Cell line authentication was performed by the Department of Human Genetics, LUMC, by using the PowerPlex® Fusion System 5C autosomal STR kit (Promega) as previously described ^48^. Both clones from two hiPSC lines had normal karyotypes, showed expression of pluripotency markers and underwent tri-lineage differentiation (Suppl. Fig. 9). Endothelial cells were generated (see below) from the following clones (donor-matched for both patients) and used in subsequent experiments: LUMC0228iCTRL03 p11 and LUMC0229iCTRL01 p11.

### Human induced pluripotent stem cells-derived endothelial cells generation and culture

Endothelial cells were differentiated from hiPSCs through mesoderm specification under defined culture conditions followed by cryopreservation until chip culture, as previously described ^49^. Cryopreserved hiPSC-derived endothelial cells were thawed in a water bath at 37 °C and transferred into a 15 mL tube containing 7 mL of Human Endothelial SFM medium (11111044; Thermo Fisher Scientific). The cells were centrifuged at 300 x g for 3 min and the pellet was resuspended in Human Endothelial SFM complete medium: Human Endothelial SFM medium supplemented with 20 ng/mL bFGF (130-093-564; Miltenyi Biotech), 30 ng/mL VEGF (130-109-396; Miltenyi Biotech), and 1% (v/v) human serum from platelet poor plasma (P2918; Sigma-Aldrich) and seeded into a T75 cell culture flask that was previously coated with 0.1% (w/v) gelatine solution for 1 h at 37 °C. The cells were expanded for 3-4 days prior to seeding into the tumor-on-chip and used only at passage 1.

### Generation and culture of ROR1-specific CAR T cells

PBMCs were isolated by density gradient centrifugation from leukocyte apheresis of two different healthy donors using a separating solution with a density of 1,077 g/mL (Pancoll human; PAN Biotech). All donors provided their written informed consent.

CD4^+^ and CD8^+^ T cells were then isolated from PBMCs by magnetic associated cell sorting (MACS) using the CD4^+^ or CD8^+^ human T cell isolation kit (130-096-533 or 130-096-495; Miltenyi Biotec), respectively. Purity of isolated T cell fractions was verified by staining with fluorophore-conjugated antibodies (all from Biolegend, San Diego, CA, USA) and 7-AAD staining solution (130-111-568; Miltenyi Biotec) for dead cell exclusion on a MACS Quant 10 analyzer (Miltenyi Biotec). The following antibodies were used: anti-human CD3 PE (clone UCHT1), anti-human CD4 APC (clone RPA-T4), anti-human CD8 FITC (clone SK1).

T cells were seeded in 48-well plates (Costar plates; Corning) in CTL medium (RPMI 1640 supplemented with 1% (v/v) Penicillin/Streptomycin, 1X GlutaMAX-I, 0.1% (v/v) 2-Mercaptoethanol [all from Thermo Fisher Scientific, Darmstadt, Germany] and 10% (v/v) pooled human serum [Bavarian Red Cross Center, Wiesentheid, Germany]) and activated using Dynabeads™ Human T-Activator CD3/CD28 Beads and 50 U/mL IL-2 (Miltenyi Biotec).

The next day (day 1), two thirds of the medium was removed from all wells and T cells were treated with 5 ng/mL polybrene (Merck, Darmstadt, Germany). Lentiviral particles (MOI=3) encoding the ROR1 CAR construct (ROR1_41BB_CD3zeta_EGFRt) were added to the cells (untransduced control: polybrene only) and centrifuged at 800 x g for 45 minutes at 32 °C. Afterwards, cells were incubated for 4 hours at 37 °C, 5% CO_2_ and 95% rH. Then, CTL medium supplemented with 50 U/mL IL-2 was added to all wells and cells were fed every other day by removing half of the medium from each well and adding CTL supplemented with 100 U/mL IL-2 (final conc. 50 U/mL IL-2). On day 7, CD3/CD28 Beads were magnetically removed and transduction efficacy was analyzed by staining with fluorophore-conjugated antibodies (all from Biolegend) and 7-AAD staining solution (Miltenyi Biotec) for dead cell exclusion on a MACS Quant 10 analyzer (Miltenyi Biotec). The following antibodies were used: anti-human EGFR Alexa Fluor® 488 (clone AY13), anti-human CD4 APC (clone RPA T4), anti-human CD8 Pacific Blue™ (clone SK1). On day 9, CAR-modified (that is, EGFRt-positive) T cells were enriched by MACS using an in-house biotinylated anti-EGFR antibody (Cetuximab, Eli Lilly and Company, Indianapolis, IN, USA) and anti-Biotin Microbeads (Miltenyi Biotec). Purity of enriched CAR modified T cells was analyzed as described above. On the following day, enriched CAR modified as well as untransduced T cells were subjected to an antigen-independent expansion protocol using irradiated CD19^+^ feeder cells as well as irradiated third-party donor PBMCs. 14 days later, purity of expanded CAR-modified T cells was analyzed as above. Then, cells were counted using a Countess Counting II FL Device (Thermo Fisher Scientific) and cryopreserved at 10 million cells/mL in Cryo SFM freezing medium (C-29910; PromoCell GmbH).

Three days before the perfusion through the chips, each subset (i.e. CD4 and CD8) of cryopreserved control T and CAR-T cells were thawed, centrifuged at 300 x g for 8 min at 8 °C, counted and seeded in 24-well cell culture plates at a concentration of 3.0 x 10^6^ cells per well. T cells were cultured for 3 days in X-VIVO^TM^ 15 medium (BE02-060F; Lonza) supplemented with 5 ng/mL of IL-15 (130-093-955; Miltenyi Biotec), 1X Glutamax and 100 U/mL penicillin-streptomycin.

### Aggregates/spheroids/organoids seeding on-chip

After chips sterilization and hydrophilization with O_2_-plasma, the channels are flushed with 70% ethanol followed by rinsing three times with PBS^−^. The chips were kept at RT until cell seeding. The cells were seeded into the chip at two to four days before T cells perfusion.

To retrieve the MDA-MB-231 aggregates from the plate, cell culture medium was firstly removed from the wells before introducing 500 µL of 2 mg/mL cold dispase II solution followed by pipetting up and down to break the gel. The plate was then incubated for 10 min at 37 °C, 5% CO_2_ and 95% rH. For the fibroblast spheroids, this gel digestion step was skipped as no gel was used for the spheroids generation. Afterwards, each well was resuspended with 1 mL of cell culture medium and the retrieved aggregates/spheroids were centrifuged down at 500 x g for 5 min. The pellet was then resuspended in dextran hydrogel (3-D Life Dextran-CD Hydrogel SG; Cellendes GmbH) supplemented with RGD peptide (09-P-001; Cellendes GmbH) with a final concentration of 0.5 mmol/L. Aggregates/spheroids from one well were used to load 3 chips.

To retrieve the organoids from the plate, gel droplets were first washed with PBS^−^ before introducing 250 µL of 2 mg/mL cold dispase II solution followed by pipetting up and down to break the gel droplet. The organoid cell suspension was then incubated for 10 min at 37 °C, 5% CO_2_ and 95% rH. Afterwards, each well was resuspended with 500 µL of advDMEM/F-12 +/+/+ and the retrieved organoids were centrifuged down at 500 x g for 5 min. The pellet was resuspended in dextran hydrogel supplemented with 0.5 mmol/L RGD peptide. Each droplet of organoids on day 10 of culture was used to load 3 chips.

In experiments that required organoids/spheroids fluorescence-labeling, organoids/spheroids pellet was resuspended in 10 µM of CellTracker^TM^ CMFDA dye solution (C7025; Invitrogen) dissolved in phenol red-free DMEM/F-12 medium and incubated for 20 min at 37 °C, 5% CO_2_ and 95% rH. The cells were then washed twice by adding phenol red-free DMEM/F-12 medium up to 10 mL and centrifuging at 500 x g for 5 min for each washing cycle. Afterwards, the pellet was resuspended in dextran hydrogel supplemented with 0.5 mmol/L RGD peptide.

Once the cells were suspended in the hydrogel, 10 µL was immediately injected into each chip via the tissue inlet (Fig 1b). The injection method resulted in approximately 3 tumor aggregates per chamber and 15 per chip, respectively (Suppl. Fig. 1c, d). In the “empty chip” condition, blank hydrogel was injected without cells. Afterwards, the inlet and outlet of the media channel were plugged using a metal wire with 0.7 mm diameter (45473; Menzanium) and the main tissue channel was flushed via negative pressure to remove any remaining cells. Then, plugs from the media channel inlet and outlet were carefully removed and inserted into the tissue channel inlet and outlet. The media channel was subsequently flushed three times with 100 µL of cell-specific culture medium to remove any deposited hydrogel that might disrupt the subsequent endothelial cells attachment and fluidic flow. The chips were then incubated at 37 °C, 5% CO_2_ and 95% rH for 30 min to allow hydrogel crosslinking.

### Endothelial cells seeding, chip culture, and characterization

MvECs were harvested by first removing the cell culture medium from the flask, rinsing with PBS^−^, followed by the insertion of 2 mL TrypLE^TM^ Select (12563-029; Gibco). The flask was incubated for 3 min at 37 °C, 5% CO_2_ and 95% rH to detach the cells. The trypsinization was stopped by adding 200 µL of FCS into the solution, followed by transferring the solution into 15 mL tube. The cells were centrifuged at 200 x g for 5 min and resuspeneded in ECGM to achieve 9 x 10^6^ cells/mL concentration for chip seeding. In case of hiPSC-ECs, the cells were harvested in a similar way with minor differences: the trypsinization process was done for 5 min at RT, and the cells were lastly centrifuged at 300 x g for 3 min at RT.

Endothelial cells were seeded into the chip on the same day of tumor cell loading and after the hydrogel in the tumor chambers was crosslinked. First, the media channel of each chip was coated with 100 µL solution containing 100 µg/mL collagen-I (FibriCol®, #5133; Advanced Biomatrix) and 20 µg/mL fibronectin (#F1141; Sigma Aldrich) for 1 h at 37 °C, 5% CO_2_ and 95% rH. The media channel was then flushed three times with ECGM. Afterwards, 5 µL of cell suspension (concentration 9 x 10^6^ cells/mL) was injected into the media channel inlet, followed by incubation for 4 h at 37 °C, 5% CO_2_ and 95% rH to allow cell attachment on the membrane. Filter pipet tips filled with 100 µL ECGM each were added into the inlet and outlet of the media channel for 24 h of static culture. The chips were cultured for 1-2 days in push mode until control T and CAR-T cells perfusion. The details of the perfusion setup are described in the section below. Prior to the T cell perfusion, random chip samples were picked for immunofluorescence staining of the CD31 expression. Briefly, CD31 antibody (anti-human, APC, REAfinity^TM^; 130-110-808; Miltenyi Biotec) [in 1:35 working concentration] and Hoechst 33342 [in 1:250 working concentration] were diluted in ECGM. 50 µL of the solution was gently introduced into the chip’s media channel followed by incubation for 30 min at 37 °C, 5% CO_2_ and 95% rH. The staining solution was rinsed gently three times with ECGM before image acquisition. Thorough characterization of the endothelium has been performed in our previous studies employing the same seeding and culture method ^10, 11^.

### T cell perfusion and chip culture

To prepare the T cells, each subset (CD4 and CD8) of the cultured control T and CAR-T cells were retrieved from the 24-well plate and pipetted through a 70 µL strainer to filter out cell clumps. Cells were centrifuged at 300 x g for 8 min at 8 °C followed by cell counting. To mimic the CAR-T cell product formulation in clinical trials, the typical 1:1 CD4^+^/CD8^+^ T cell ratio was selected throughout our experiments ^25, 26^, as it has shown a superior antitumor reactivity in an in vivo study and their cooperation is essential to exert potent and long-lasting antitumor activity ^50–52^. Prior to the perfusion, both subsets were mixed and the concentration was set to 0.5 x 10^6^ cells/mL. The concentration was set to 100,000 cells/mL and 1,000,000 cells/mL only in preliminary experiments to determine the optimal perfused T cell concentration (Suppl. Fig. 3b). In experiments that required fluorescence-labeling of T cells, T cell pellet was resuspended in 2 µM of CellTracker^TM^ Deep Red dye solution (C34565; Invitrogen) dissolved in phenol red- and serum-free DMEM/F-12 medium and incubated for 15 min at 37 °C, 5% CO_2_ and 95% rH. The cells were then washed twice by adding fully supplemented X-VIVO^TM^ 15 medium up to 10 mL and centrifuging at 300 x g for 8 min at 8 °C for each washing cycle. Afterwards, the final cell pellet was resuspended in ECGM basal medium (C-22210; PromoCell GmbH) supplemented with 2% FCS, 12 µg/mL ECGS, 90 µg/mL heparin, 0.1 ng/mL EGF, 1 ng/mL bFGF (all from the Growth Medium SupplementPack C-39210; PromoCell GmbH) – this will be referred to as “co-culture medium” in the following.

Day of control T and CAR-T cell perfusion was defined as day 0 for all experiments. Starting from day 0, the chips were connected to constant media perfusion via an external syringe pumping system (LA-190, Landgraf Laborsysteme HLL GmbH, Langenhagen, Germany). The chips were connected to the syringe pump using blunt 21 GA stainless steel needles (made from the dispensing needles by removing the plastic hub after dissolving the glue overnight in a 70% ethanol solution) connected to Tygon tubings (0.51 mm inner diameter, Tygon ND 100-80 Medical Tubing, Saint-Gobain Performance Plastics Pampus GmbH, Willich, Germany), 21 GA stainless steel plastic hub dispensing needles (e.g., KDS2112P, Weller Tools GmbH, Besigheim, Germany) and Luer Lock style syringes. For (CAR-)T cell perfusion through the media channel, the inlet of the channel was equipped with a 100 µL-pipette tip reservoir holding the 400 µL of T cell suspension (200,000 T cells/chip). The syringe pump was set to the withdraw mode at 100 µL/h to ensure steady flow before introducing the (CAR-)T cells, thereby preventing them to precipitate on the inlet area. Once the (CAR-)T cells were suspended in the pipette tip reservoir, the flow rate was set to 20 µL/h and the whole setup was transferred into an incubator at 37 °C, 5% CO_2_ and 95% rH. After 20 h, the chip effluents and plate control supernatants were collected for further analysis and the perfusion setup was changed to push mode where co-culture medium was dispensed from the syringes. The chips further linearly perfused in this mode at 20 µL/h flow rate until up to day 8. Medium in the syringes was refilled after 3 days of perfusion. Before chip perfusion, medium is warmed-up to 37 °C and simultaneously degassed (vacuum source −70 kPa) and filter-sterilized using a Steriflip® conical filter unit for 20 min. Effluent samples from each chip were collected every one or two days, centrifuged at 200 x g for 5 min to remove perfused cells, then at 10,000 x g for 10 min to remove debris and stored at −80 °C until further analysis.

### Dasatinib treatment

To model CRS intervention, dasatinib (S1021; Selleckchem) with a final concentration of 50 nM was supplemented in the co-culture medium and perfused through the media channel of the chips either from day 0-1, 0-8, or 1-8. This concentration was found to be potent to render CD19-CAR-T cells temporarily inactive ^39^. In a preliminary experiment, CAR-T cells were cultured either with or without MDA-MB-231 cells (effector-to-target cell ratio, 10:1) for 24 h, with or without dasatinib supplementation (50 nM final concentration) in the co-culture medium. In the co-culture condition, MDA-MB-231 cells were seeded as a monolayer and CAR-T cells (1:1 ratio of CD8^+^ and CD4^+^) suspension was added into the well. CAR-T cells were collected on the next day for flow cytometry staining with anti-human CD69 (early activation marker), anti-human CD25 (late activation marker). Viability was assessed also during flow cytometry by adding Zombie aqua dye (423102; BioLegend) into the staining solution (see “Flow cytometry staining and analysis” method section).

### Allogeneic / isogenic endothelial cells – (CAR-)T cells co-culture for cytokine assay

To demonstrate the differences of allogeneic and isogenic setup in terms of cytokine release by the T cells, we selected five mvECs donors for 24 h co-culture with T cells. HiPSC-ECs was used in the isogenic setup. Briefly, endothelial cells were plated (10,000 cells/well, 96-well plate format) and cultured overnight. On the next day, 200,000 control T or CAR-T cells (1:1 ratio of CD8^+^ and CD4^+^) were added into each well (effector-to-target cell ratio, 20:1). Cells were co-cultured overnight and co-culture medium was used: see method section “T cell perfusion and chip culture” for the composition. Supernatants were collected and stored at −80 °C until cytokine analysis (see “Evaluation of chip effluents and plate control supernatants” method section).

### Cytokine quantification assay

Cytokines from undiluted samples from chips/plates were analyzed by fluorescent bead-based multiplex sandwich immunoassays (Legendplex™ Human CD8/NK Mix and Match Subpanel; BioLegend) containing capture beads targeting IL-2, IL-6, IL-10, TNF-α, IFN-γ and granzyme B and read by flow cytometry (BD LSR II or BD LSRFortessa™, BD Biosciences) according to manufacturer’s protocol. Flow cytometry data files were manually gated using LEGENDplex Cloud-Based Data Analysis Software Suite (BioLegend) to find the optimal differentiation between capture bead populations and subsequently applied to all datasets.

### Flow cytometry staining and analysis

For the analysis of the expression of extracellular markers, the following antibodies were used depending on the experiment: anti-human ROR1 APC (130-118-015; Miltenyi Biotec), anti-human CD31 FITC (130-110-806; Miltenyi Biotec), anti-human CD3 PE/Cyanine7 (317334; BioLegend), anti-human CD4 APC/Fire750 (357426; BioLegend), anti-human CD8a PerCP (300922; BioLegend), anti-human CD69 FITC (310904; BioLegend) and anti-human CD25 BV421 (302630; BioLegend). Detailed information on the antibodies is summarized in Supplementary Table 2. 96-well V-bottom cell culture plate was used for staining and flow cytometry reading. For staining, cells in each well were firstly washed with flow cytometry staining buffer: 3% FCS with 2 mM EDTA (15575020; Invitrogen) in PBS^−^. Cells were then incubated in a blocking solution containing Human BD Fc Block (564220; BD Biosciences) diluted 1:25 (v/v) in PBS^−^ for 15 min at 4 °C. Afterwards, cells were incubated for 30 min at 4° C in the antibody-mixture solution specific for each experiment. Zombie aqua dye (423102; BioLegend) was added 1:200 in the antibody-mixture solution to differentiate the viable cells during analysis. Cells were then washed twice with the staining buffer, fixed with ROTIHistofix 4% (P087.6; Carl Roth GmbH) for 10 min at RT, then washed and resuspended in the staining buffer. The samples were analyzed directly using flow cytometry (BD LSR II or BD LSRFortessa™, BD Biosciences) or kept stored at 4 °C for maximum overnight until analysis. Autofluorescence and isotype controls were included in every staining procedure. For the analysis, FSC-A and SSC-A properties were used to identify cells, cell doublets were excluded based on FSC-H vs. FSC-A. Dead cells were excluded for analysis. Primary cancer cells, cancer cell lines and endothelial cells were analyzed with established ROR1 marker by using unstained cells as cut-off. Same strategy was applied for CD31 staining on hiPSC-derived endothelial cells. For T cell analyis, T cell subpopulations were gated according to established markers and analyzed for activation marker expression by comparing activation marker signal in untreated vs. treated T cells.

### Image acquisition and analysis

For the timepoint live imaging of T cell infiltration, GFP fluorescence intensity of the MDA-MB-231 tumor aggregates and tumor aggregate area, an epifluorescence microscope (Axio Observer 7, Carl Zeiss) was used and the focal plane was set to focus on the aggragate/spheroid/PDO. The microscope chamber temperature was set to 37 °C during all live cell imaging acquisitions of the chips. Analyses of T cell infiltration, GFP fluorescence intensity of the MDA-MB-231 tumor aggregates and tumor aggregate area were conducted using ImageJ-Fiji software ^53^. For T cell intensity measurements, the histogram of the fluorescent images was adjusted to remove the background signal, followed by quantification of the mean gray value of the CellTracker^TM^ Deep Red-labeled (CAR-)T cells within the region of interest (ROI) of each tumor aggregate/fibroblast spheroid/PDO. Tumor aggregates/fibroblasts spheroids/PDOs smaller than 50 µm in diameter were excluded from the analysis. Tumor aggregate/fibroblasts spheroid/PDO ROI was defined by automated thresholding of the GFP signal (for MDA-MB-231 aggregates) or the CellTracker^TM^ CMFDA signal (for fibroblasts spheroids and PDOs). Area of each MDA-MB-231 aggregate was measured after thresholding to analyze tumor aggregate growth upon control T or CAR-T cell treatment, and aggregate area from day > 0 was compared with their initial area before control T or CAR-T cell perfusion. For GFP fluorescence intensity analysis of the tumor aggregate, the mean gray value within each tumor aggregate ROI was measured and compared with their initial mean gray value on day 0. To quantify T cells in the whole tumor chamber, a confocal Laser-Scanning-Microscope (LSM 880, Carl Zeiss) was used. Here, confocal z-stacks imaging data were used to run the Spot Detection function in Imaris 9.5 software (Oxford Instruments). The estimated XY and Z diameter of T cells was set to 5 µm and 10 µm, respectively. For all measurements, image data from a minimum of 3 chips were pooled for each condition for further statistical analysis. For the quantification, three chambers were selected per chip (the first, third, and sixth). If no cell was present in one of the chambers, the second and/or the fourth chamber was included for the analysis. To measure MDA-MB-231 aggregates diameter after the aggregate generation method using the agarose microwells-based approach, a randomly selected area in the well (three wells in total) was imaged and further processed for automated thresholding and subsequent diameter measurement in ImageJ-Fiji to check for the diameter variability.

### Statistics and reproducibility

In all experiments, every chip is considered an independent biological replicate. Chips were excluded from the analysis if technical issues with the microfluidic setup occurred. The number of biological replicates is described in each figure caption. For imaging-based quantification, n denotes the number of aggregates/spheroids/PDOs. For the effluent analysis, n denotes the number of chip replicates. Statistical analysis was performed using GraphPad Prism 9.2.0 software, and graphs are represented as average values ± SEM unless indicated otherwise. Detailed statistics are indicated in each figure legend.

## ACKNOWLEDGEMENTS

We thank E. Bras, G. Bae, G. Sharma, and A. S. Schlemmer for assisting the chip fabrication, C. Önder for assisting the flow cytometry of the organoids, the LUMC human iPSC Hotel for the generation and initial characterization of hiPSC lines from PBMCs, the Flow Cytometry Core Facility Tal at the University of Tübingen, and the imSAVAR consortium.

## AUTHORS CONTRIBUTIONS

T.I.M. performed the chip design and fabrication, as well as cell culture and chip experiments with assistance from C.T., L.L and M.C. C.T. and L.L. performed the well-plate assays of preliminary 2D experiments. C.T. and A.K. performed the flow cytometry and analyzed the data. A.K. provided the organoids. M.A. and M.H. provided the (CAR-)T cells and intellectual expertise on the CAR-T cell engineering. F.E.v.d.H. and V.O. provided the hiPSCs and hiPSC-derived endothelial cells and performed their characterization. T.I.M., M.C., M.A. and P.L. designed the study. T.I.M. performed the imaging and image data analysis. T.I.M. analyzed the data from the chip experiments. T.I.M. wrote the manuscript with support from P.L., M.C., C.T., M.A., A.K., V.O., M.A., and M.H. This work was jointly supervised by P.L. and M.H. who share senior authorship. All authors have read the manuscript, provided input, and agree with its submission.

## REFERENCES

1. June, C. H., O’Connor, R. S., Kawalekar, O. U., Ghassemi, S. & Milone, M. C. CAR T cell immunotherapy for human cancer. Science 359, 1361–1365 (2018).

2. Hou, A. J., Chen, L. C. & Chen, Y. Y. Navigating CAR-T cells through the solid-tumour microenvironment. Nat Rev Drug Discov 20, 531–550 (2021).

3. Morris, E. C., Neelapu, S. S., Giavridis, T. & Sadelain, M. Cytokine release syndrome and associated neurotoxicity in cancer immunotherapy. Nat Rev Immunol 22, 85–96 (2022).

4. Xiao, X. et al. Mechanisms of cytokine release syndrome and neurotoxicity of CAR T-cell therapy and associated prevention and management strategies. J Exp Clin Cancer Res 40, 367 (2021).

5. Wei, J. et al. The model of cytokine release syndrome in CAR T-cell treatment for B-cell non-Hodgkin lymphoma. Sig Transduct Target Ther 5, 134 (2020).

6. Sterner, R. C. & Sterner, R. M. CAR-T cell therapy: current limitations and potential strategies. Blood Cancer J. 11, 69 (2021).

7. Donnadieu, E. et al. Time to evolve: predicting engineered T cell-associated toxicity with next-generation models. J Immunother Cancer 10, e003486 (2022).

8. Guedan, S. et al. Time 2EVOLVE: predicting efficacy of engineered T-cells – how far is the bench from the bedside? J Immunother Cancer 10, e003487 (2022).

9. Maulana, T. I. et al. Immunocompetent cancer-on-chip models to assess immuno-oncology therapy. Adv Drug Deliv Rev 173, 281–305 (2021).

10. Cipriano, M. et al. Human immunocompetent choroid-on-chip: a novel tool for studying ocular effects of biological drugs. Commun Biol 5, 52 (2022).

11. Rogal, J. et al. Autologous Human Immunocompetent White Adipose Tissue-on-Chip. Advanced Science 9, 2104451 (2022).

12. Rafiq, S., Hackett, C. S. & Brentjens, R. J. Engineering strategies to overcome the current roadblocks in CAR T cell therapy. Nat Rev Clin Oncol 17, 147–167 (2020).

13. Hudecek, M. et al. The B-cell tumor–associated antigen ROR1 can be targeted with T cells modified to express a ROR1-specific chimeric antigen receptor. Blood 116, 4532–4541 (2010).

14. Zhang, S. et al. The Onco-Embryonic Antigen ROR1 Is Expressed by a Variety of Human Cancers. The American Journal of Pathology 181, 1903–1910 (2012).

15. Broome, H. E., Rassenti, L. Z., Wang, H.-Y., Meyer, L. M. & Kipps, T. J. ROR1 is expressed on hematogones (non-neoplastic human B-lymphocyte precursors) and a minority of precursor-B acute lymphoblastic leukemia. Leukemia Research 35, 1390–1394 (2011).

16. Joyce, J. A. & Fearon, D. T. T cell exclusion, immune privilege, and the tumor microenvironment. Science 348, 74–80 (2015).

17. Jiang, P. et al. Signatures of T cell dysfunction and exclusion predict cancer immunotherapy response. Nat Med 24, 1550–1558 (2018).

18. Dekkers, J. F. et al. Uncovering the mode of action of engineered T cells in patient cancer organoids. Nat Biotechnol (2022) doi:10.1038/s41587-022-01397-w.

19. Yu, L. et al. Patient-derived organoids of bladder cancer recapitulate antigen expression profiles and serve as a personal evaluation model for CAR-T cells in vitro. Clin Transl Immunol 10, (2021).

20. Jacob, F., Ming, G. & Song, H. Generation and biobanking of patient-derived glioblastoma organoids and their application in CAR T cell testing. Nat Protoc 15, 4000–4033 (2020).

21. Arenas, E. J. et al. Acquired cancer cell resistance to T cell bispecific antibodies and CAR T targeting HER2 through JAK2 down-modulation. Nat Commun 12, 1237 (2021).

22. Schnalzger, T. E. et al. 3D model for CAR-mediated cytotoxicity using patient-derived colorectal cancer organoids. EMBO J 38, (2019).

23. Halloin, C. et al. Continuous WNT Control Enables Advanced hPSC Cardiac Processing and Prognostic Surface Marker Identification in Chemically Defined Suspension Culture. Stem Cell Reports 13, 366–379 (2019).

24. Mazein, A. et al. Using interactive platforms to encode, manage and explore immune-related adverse outcome pathways. http://biorxiv.org/lookup/doi/10.1101/2023.03.21.533620 (2023) doi:10.1101/2023.03.21.533620.

25. Turtle, C. J. et al. Immunotherapy of non-Hodgkin’s lymphoma with a defined ratio of CD8 ^+^ and CD4 ^+^ CD19-specific chimeric antigen receptor–modified T cells. Sci. Transl. Med. 8, (2016).

26. Turtle, C. J. et al. CD19 CAR–T cells of defined CD4+:CD8+ composition in adult B cell ALL patients. Journal of Clinical Investigation 126, 2123–2138 (2016).

27. Teachey, D. T. et al. Identification of Predictive Biomarkers for Cytokine Release Syndrome after Chimeric Antigen Receptor T-cell Therapy for Acute Lymphoblastic Leukemia. Cancer Discovery 6, 664–679 (2016).

28. Tedesco, V. E. & Mohan, C. Biomarkers for Predicting Cytokine Release Syndrome following CD19-Targeted CAR T Cell Therapy. The Journal of Immunology 206, 1561–1568 (2021).

29. Hay, K. A. et al. Kinetics and biomarkers of severe cytokine release syndrome after CD19 chimeric antigen receptor–modified T-cell therapy. Blood 130, 2295–2306 (2017).

30. Norelli, M. et al. Monocyte-derived IL-1 and IL-6 are differentially required for cytokine-release syndrome and neurotoxicity due to CAR T cells. Nat Med 24, 739–748 (2018).

31. Maude, S. L. et al. Chimeric Antigen Receptor T Cells for Sustained Remissions in Leukemia. N Engl J Med 371, 1507–1517 (2014).

32. Lee, D. W. et al. T cells expressing CD19 chimeric antigen receptors for acute lymphoblastic leukaemia in children and young adults: a phase 1 dose-escalation trial. The Lancet 385, 517–528 (2015).

33. Grupp, S. A. et al. Chimeric antigen receptor-modified T cells for acute lymphoid leukemia. N Engl J Med 368, 1509–1518 (2013).

34. Neelapu, S. S. et al. Chimeric antigen receptor T-cell therapy — assessment and management of toxicities. Nat Rev Clin Oncol 15, 47–62 (2018).

35. Neelapu, S. S. Managing the toxicities of CAR T-cell therapy. Hematological Oncology 37, 48– 52 (2019).

36. Neelapu, S. S. et al. Axicabtagene Ciloleucel CAR T-Cell Therapy in Refractory Large B-Cell Lymphoma. N Engl J Med 377, 2531–2544 (2017).

37. Maude, S. L. et al. Tisagenlecleucel in Children and Young Adults with B-Cell Lymphoblastic Leukemia. N Engl J Med 378, 439–448 (2018).

38. Giavridis, T. et al. CAR T cell–induced cytokine release syndrome is mediated by macrophages and abated by IL-1 blockade. Nat Med 24, 731–738 (2018).

39. Mestermann, K. et al. The tyrosine kinase inhibitor dasatinib acts as a pharmacologic on/off switch for CAR T cells. Sci. Transl. Med. 11, eaau5907 (2019).

40. Rogal, J. et al. WAT-on-a-chip integrating human mature white adipocytes for mechanistic research and pharmaceutical applications. Sci Rep 10, 6666 (2020).

41. Xia, Y. & Whitesides, G. M. SOFT LITHOGRAPHY. Annu. Rev. Mater. Sci. 28, 153–184 (1998).

42. Dahlmann, J. et al. The use of agarose microwells for scalable embryoid body formation and cardiac differentiation of human and murine pluripotent stem cells. Biomaterials 34, 2463–2471 (2013).

43. Miyoshi, H. & Stappenbeck, T. S. In vitro expansion and genetic modification of gastrointestinal stem cells in spheroid culture. Nat Protoc 8, 2471–2482 (2013).

44. Okita, K. et al. A more efficient method to generate integration-free human iPS cells. Nat Methods 8, 409–412 (2011).

45. Bouma, M. J., et al. Differentiation-Defective Human Induced Pluripotent Stem Cells Reveal Strengths and Limitations of the Teratoma Assay and In Vitro Pluripotency Assays. Stem Cell Reports 8, 1340–1353 (2017).

46. Dambrot, C. et al. Polycistronic lentivirus induced pluripotent stem cells from skin biopsies after long term storage, blood outgrowth endothelial cells and cells from milk teeth. Differentiation 85, 101–109 (2013).

47. Freund, C., Davis, R. P., Gkatzis, K., Ward-van Oostwaard, D. & Mummery, C. L. The first reported generation of human induced pluripotent stem cells (iPS cells) and iPS cell-derived cardiomyocytes in the Netherlands. Neth Heart J 18, 51–54 (2010).

48. Westen, A. A. et al. Comparing six commercial autosomal STR kits in a large Dutch population sample. Forensic Science International: Genetics 10, 55–63 (2014).

49. Orlova, V. V. et al. Generation, expansion and functional analysis of endothelial cells and pericytes derived from human pluripotent stem cells. Nat Protoc 9, 1514–1531 (2014).

50. Sommermeyer, D. et al. Chimeric antigen receptor-modified T cells derived from defined CD8+ and CD4+ subsets confer superior antitumor reactivity in vivo. Leukemia 30, 492–500 (2016).

51. Ostroumov, D., Fekete-Drimusz, N., Saborowski, M., Kühnel, F. & Woller, N. CD4 and CD8 T lymphocyte interplay in controlling tumor growth. Cell. Mol. Life Sci. 75, 689–713 (2018).

52. Raskov, H., Orhan, A., Christensen, J. P. & Gögenur, I. Cytotoxic CD8+ T cells in cancer and cancer immunotherapy. Br J Cancer 124, 359–367 (2021).

53. Schindelin, J., et al. Fiji: an open-source platform for biological-image analysis. Nat Methods 9, 676–682 (2012).

