## Supplementary Materials for "Solid tumor-on-chip model for efficacy and safety assessment of CAR-T cell therapy"

Prof. Dr. rer. nat. Peter Loskill, Department of Microphysiological Systems, Eberhard-Karls-University-Tübingen, Österbergstrasse 3, 72074 Tübingen, Germany.

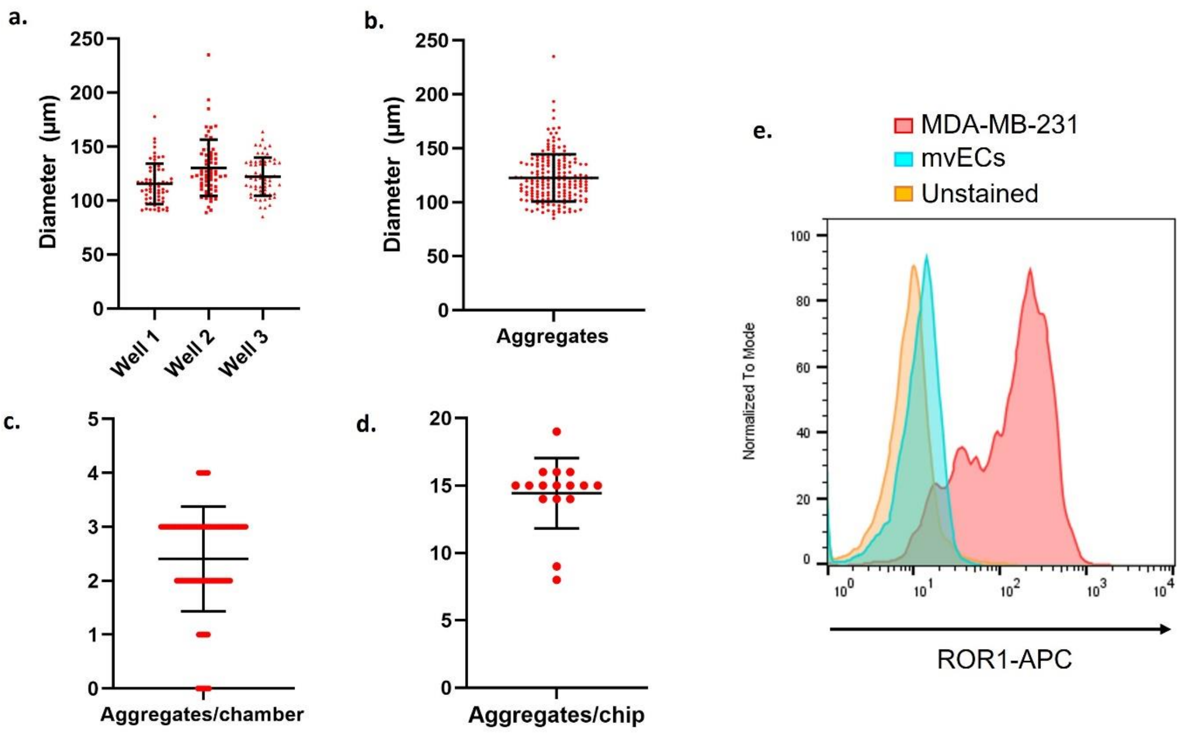


**Supplementary Figure 1: Tumor aggregates characterization and ROR1 expression.**

**a**, Diameter of the MDA-MB-231 aggregates on day 3 after being generated using the agarose microwells-based approach. *n* = 62-66 aggregates. Data depicted are mean ± SD. **b**, Pooled data points from (a.); *n* = 190 aggregates. **c**, Quantification of MDA-MB-231 aggregates per chamber after chip loading; *n* = 96 chambers from 16 chips. **d**, Quantification of MDA-MB-231 aggregates per chip after loading; *n* = 16 chips. **e**, Flow cytometry histogram plots showing ROR1 expression in the MDA-MB-231 cells (red), mvECs (cyan) compared with the unstained control (orange).

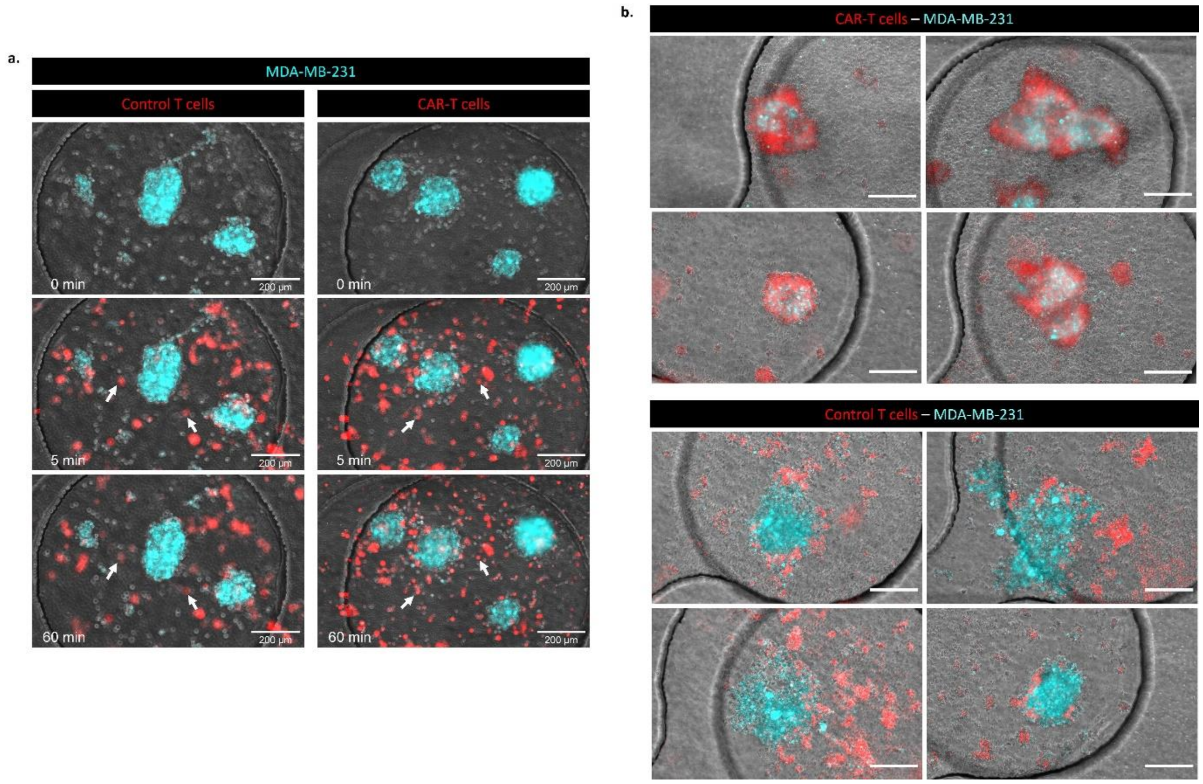


**Supplementary Figure 2: (CAR-)T cell migration towards and infiltration into the MDA-MB-231 tumor aggregates.**

**a**, Representative images during the first 1 h of control T or CAR-T cell perfusion through the tumor-on-chip containing MDA-MB-231 aggregates. The focal plane was set on the aggregates throughout the entire imaging acquisition time. The arrow indicates the T cells that either stayed out of focus and remained in the media channel (control T cell condition) or went towards focus and were increasingly found to be on the same focal plane as the aggregates (CAR-T cell condition). The time point of imaging is indicated at the bottom left of each image. Scalebar: 200 µm. **b**, Representative images from four different chips of infiltrating CAR-T cells (top four images) and excluded control T cells from the MDA-MB-231 tumor aggregates (bottom four images) on day 8. Scalebar: 200 µm.

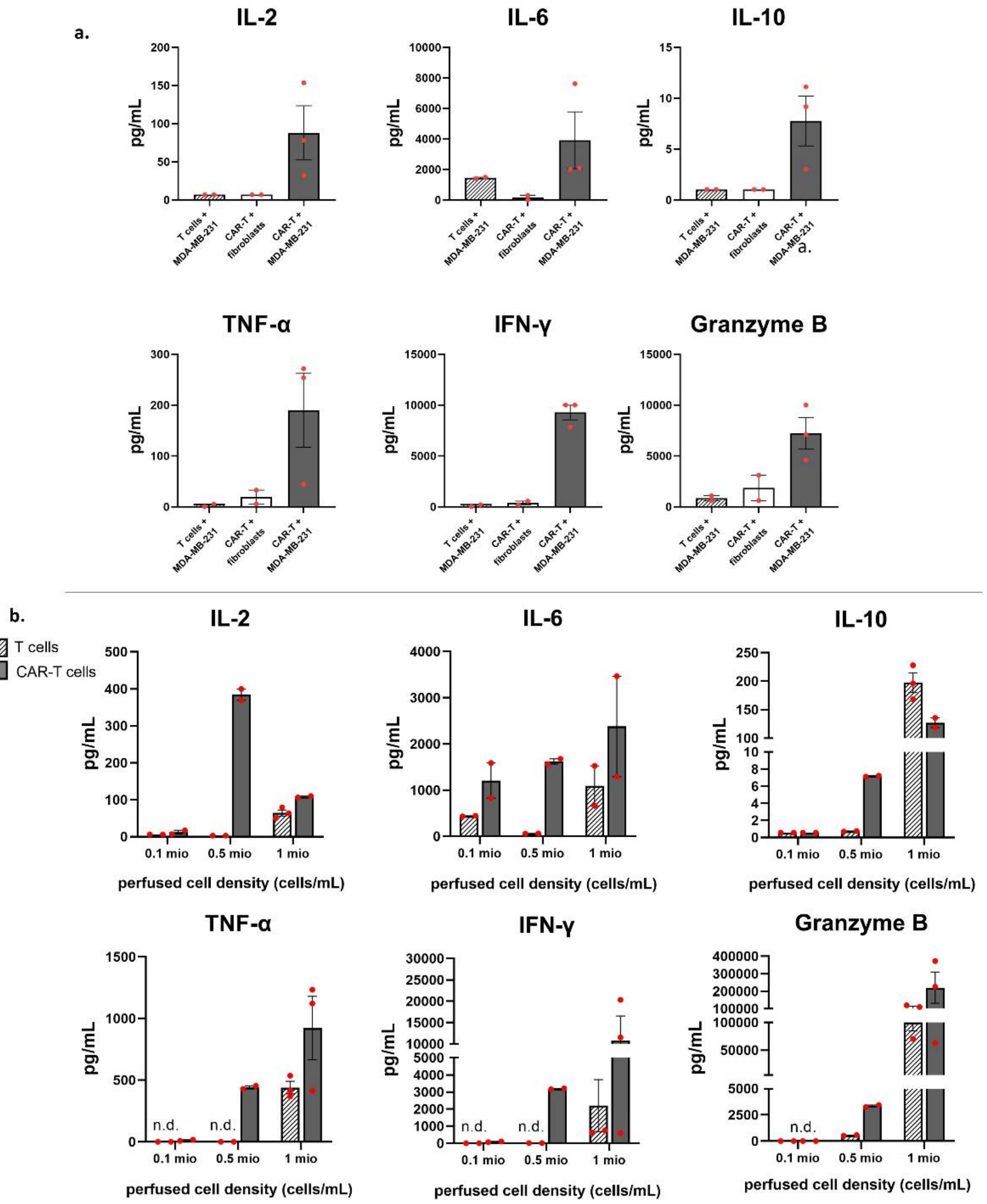


**Supplementary Figure 3: Impact of perfused (CAR-)T cell concentration and target cells on cytokine levels on day 1 post-perfusion.**

**a**, Quantification of the levels of cytokines IL-2, IL-6, IL-10, TNF-α, IFN-γ and granzyme B in the effluents of the chips after 20 h of (control) T or CAR-T cells perfusion through chips containing $ROR1^-$ fibroblasts spheroids. $ROR1^+$ MDA-MB-231 aggregates + CAR-T cells condition was used as positive control, and $ROR1^+$ MDA-MB-231 aggregates + (control) T cells condition was used as negative control. (CAR-)T cells were perfused at 500,000 cells/mL; *n* = 2-3 chips, as indicated by the red dot. Data are depicted as mean with ± SEM. **b**, Quantification of the levels of cytokines IL-2, IL-6, IL-10, TNF-α, IFN-γ and granzyme B in the effluents of the chips after 20 h of (control) T or CAR-T cells perfusion through tumor-on-chips containing MDA-MB-231 aggregates. The concentration of the perfused cells was varied: 100,000, 500,000, and 1,000,000 cells/mL; *n* = 2-3 chips, indicated by the red dot. Data are depicted as mean with ± SEM. n.d.: not detected, values below the detection limit.

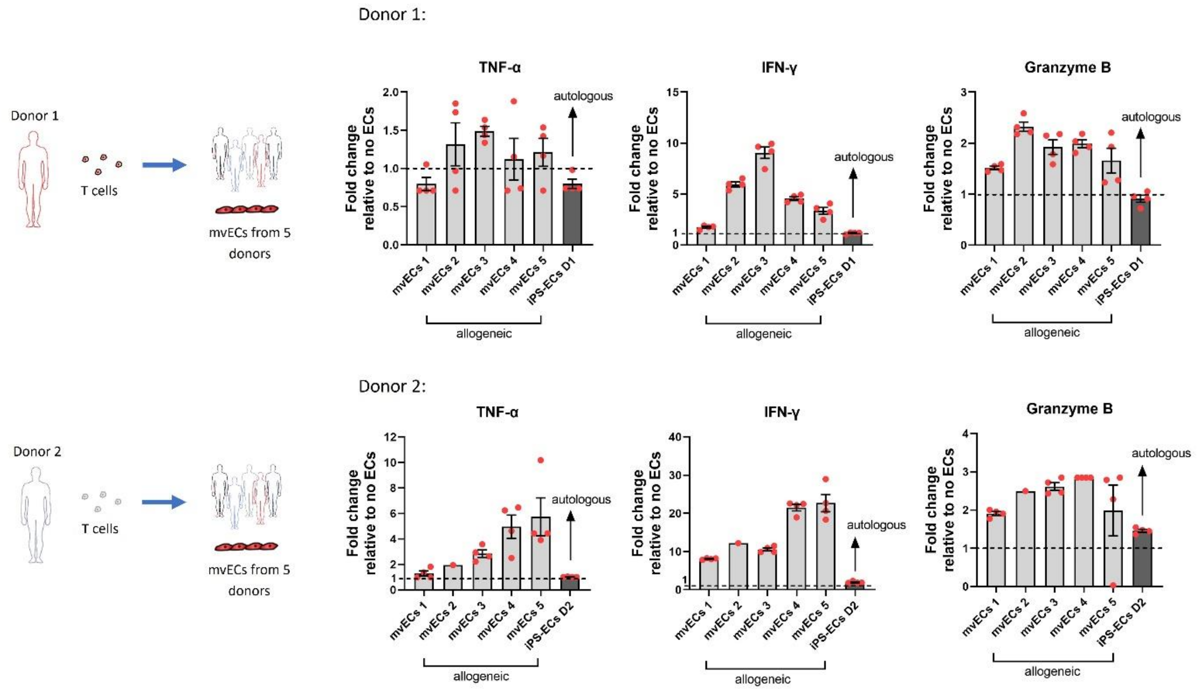


**Supplementary Figure 4: Cytokine secretion upon T cells – endothelial cells co-culture either in allogeneic or isogenic/autologous setting.**

Fold change cytokine values of TNF-α, IFN-γ and granzyme B from T cells sourced from two different donors co-cultured for 24 h either in allogeneic or isogenic/autologous setting relative to the culture without endothelial cells. In the allogeneic setting, T cells were co-cultured with mvECs from 5 different donors. T cells were co-cultured with iPSC-ECs in autologous setting, which were differentiated from iPSCs previously reprogrammed from PBMCs of the same T cell donor. Each dot represents a data point from one cell culture well. Data are shown as mean with ± SEM; *n* = 1-4.

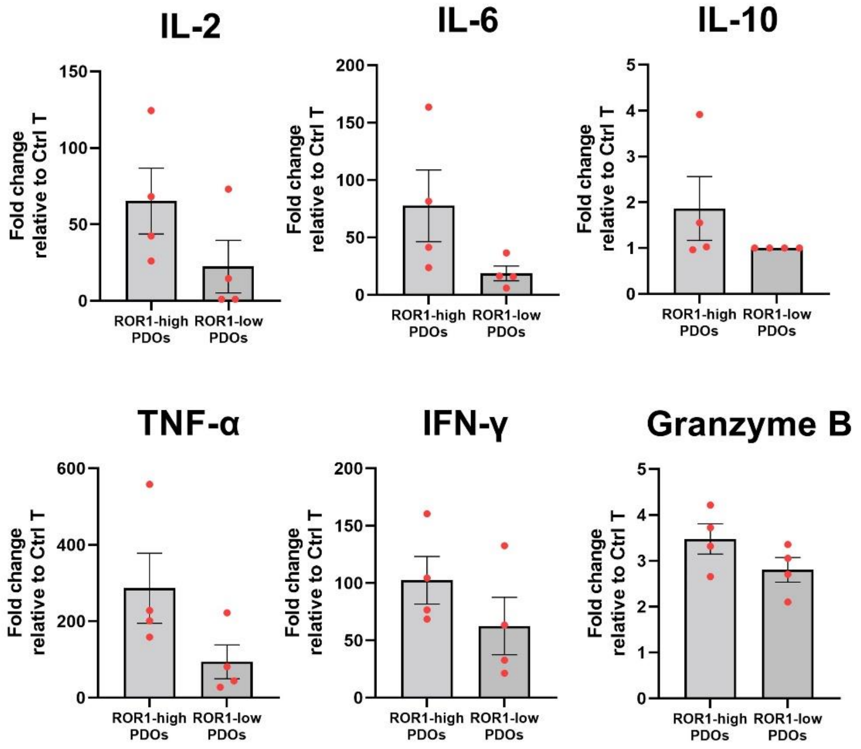


**Supplementary Figure 5: Cytokine secretion of CAR-T cells against two different PDOs expressing different levels of ROR1 antigen.**

Fold change cytokine values from day 1 obtained from the tumor-on-chips containing either ROR1-high- or ROR1-low-expressing PDOs perfused with CAR-T cells, relative to the respective control T cells condition, showing a trend of higher fold change in ROR1-high PDOs condition. Each dot represents effluent data collected from one chip. Data are shown as mean with ± SEM; *n* = 4.

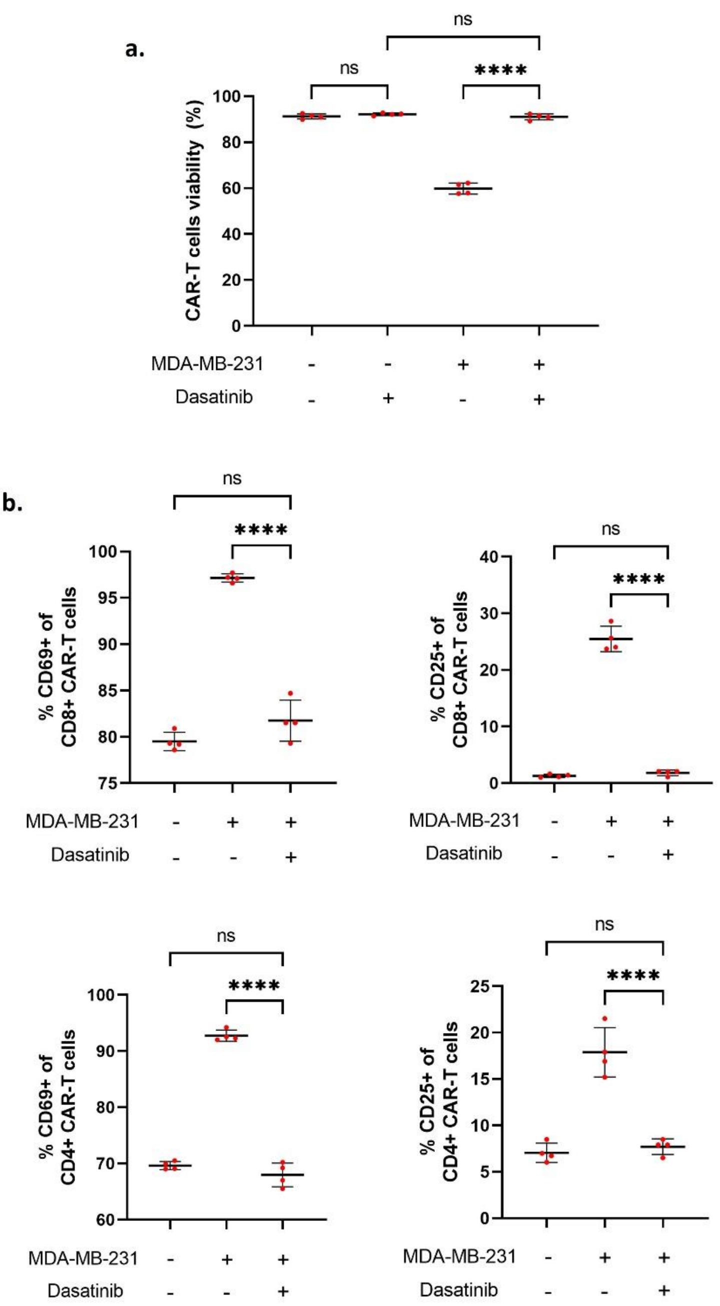


**Supplementary Figure 6: CAR-T cells viability and activation status upon dasatinib treatment.**
**a**, CAR-T cells viability after 24 h of dasatinib (50 nM) treatment either in the presence of MDA-MB-231 cells (effector-to-target cell ratio, 10:1) or without, cultured in a well plate format. **b**, Percentage of CD69$^+$ (early activation) and CD25$^+$ (late activation) CAR-T cells within the CD8$^+$ and CD4$^+$ subset after 24 h of dasatinib (50 nM) treatment either in the presence of MDA-MB-231 cells (effector-to-target cell ratio, 10:1) or without, cultured in a well plate format. Data were acquired from flow cytometry analysis for both (a.) and (b.); *n* = 4 wells. Data are depicted as mean with ± SEM. Each red dot represents one well. ns: not significant, ****p<0.0001; Tukey's multiple comparisons test.

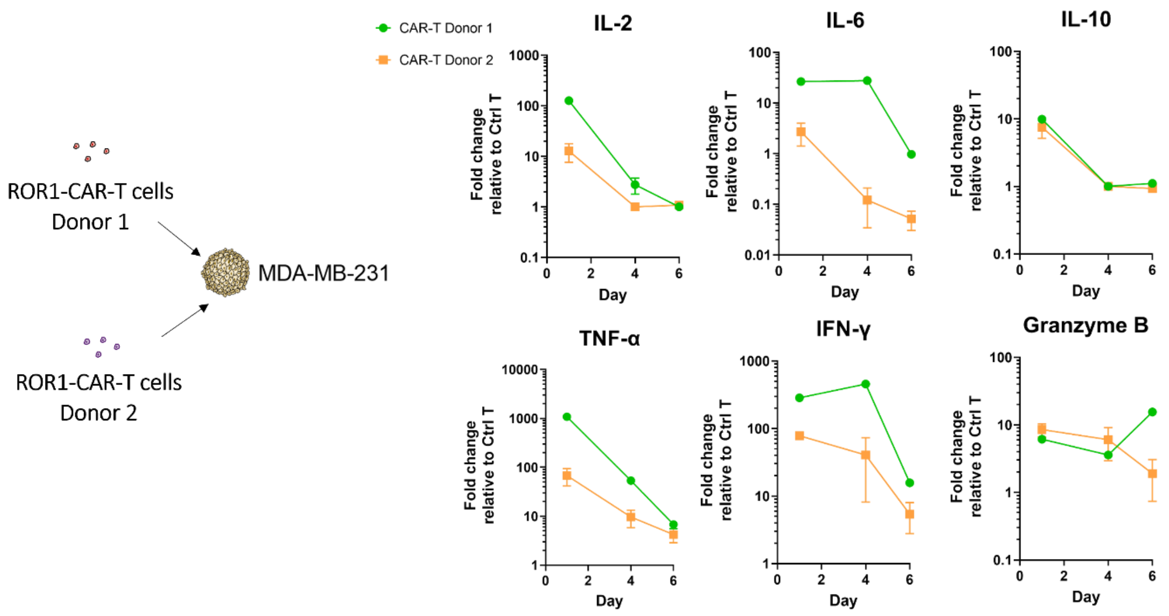


**Supplementary Figure 7: Cytokine secretion kinetics of CAR-T cells from two different donors after tumor-on-chips perfusion.**

Fold change cytokine values from day 1, 4 and 6 obtained from the tumor-on-chips containing mvECs barrier and MDA-MB-231 aggregates perfused with CAR-T cells, relative to the respective control T cells condition. CAR-T cells were sourced from two different healthy donors and were perfused through the tumor-on-chips in two independent experiments. Each dot represents effluent data collected from one chip. Data are shown as mean with ± SEM; *n* = 2-3.

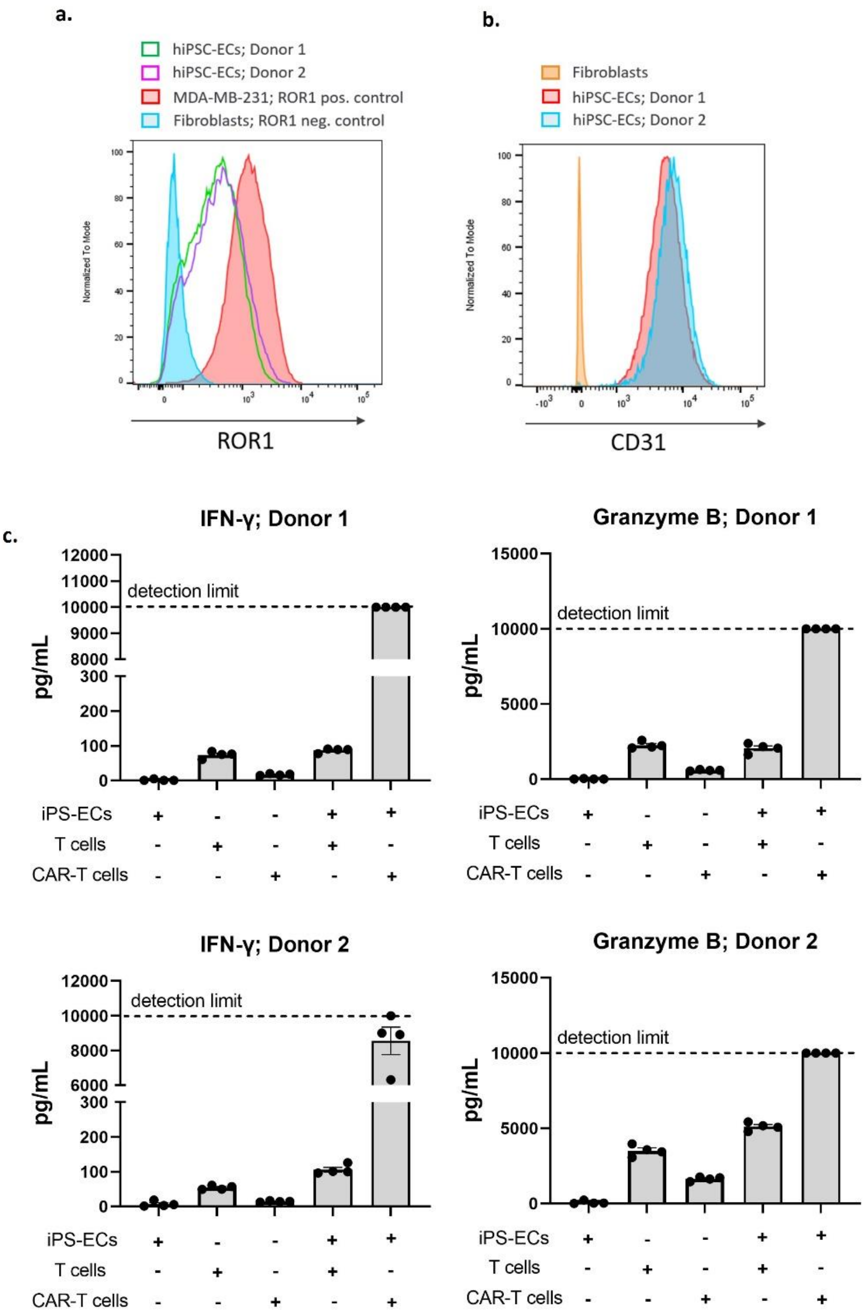


**Supplementary Figure 8: HiPSCs characterization for ROR1 and CD31 expression as well as effects of their co-culture with (CAR-)T cells on cytokine secretion.**

**a**, Flow cytometry histogram plots showing ROR1 expression levels in the hiPSC-ECs from two different donors, as well as MDA-MB-231 and fibroblasts as the positive and negative control, respectively. **b**, Flow cytometry histogram plots showing CD31 expression of hiPSC-ECs from two different donors used in the experiments integrating PDOs. Fibroblasts were used as the negative control. **c**, Quantification of the IFN-γ and granzyme B in the effluents of a plate culture experiment after 24 h of (control) T or CAR-T cells co-culture with or without autologous hiPSC-ECs. Single cultures were used as the control conditions. Cells derived from two donors were tested; *n* = 4. Data are depicted as mean with ± SEM. Each red dot represents the cytokine value from one well.

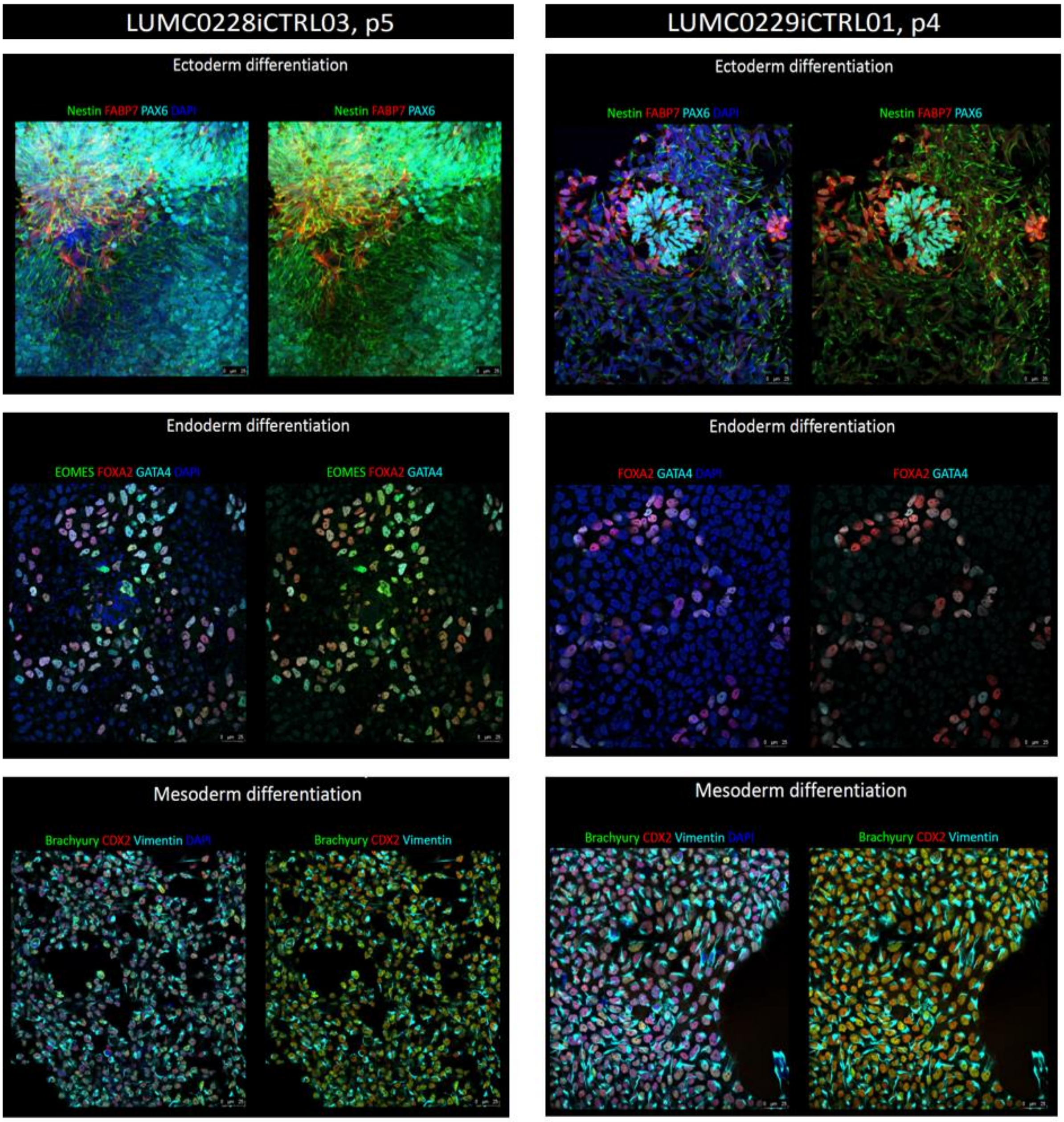


**Supplementary Figure 9: Characterization of hiPSCs' capability for tri-lineage differentiation.**
Two hiPSCs lines generated from PBMCs from healthy donors differentiated to precursors of ectoderm, endoderm, and mesoderm and stained with specific antibodies: Nestin, FABP7, PAX6 (ectoderm), EOMES, FOXA2, GATA4 (endoderm), Brachury, CDX2, Vimentin (Mesoderm). DAPI visualizes the nuclei. Scale bar 25 µm.

**Supplementary Table 1: Patient data for the PDOs used in the study.**

| Line | Age | Primary tumor | | Metastasis | | Treatment | | PDO |
|---|---|---|---|---|---|---|---|---|
| | | Type | Receptor status | Type | Receptor status | Drainage | Prior therapy | Receptor status |
| (#10) PDO-ROR1-high | 63 | IBC (2020) | ER-/PR-/HER2- | Lymphogenic and cutaneous metastasis | ER-/PR-/HER2- | Pleural effusion | Carboplatin + Paclitaxel + Atezolizumab | ER-/PR-/HER2- |
| (#43) PDO-ROR1-low | 52 | NST (2019) | ER+/PR+/HER2- | Hepatic, osseus, and pulmonary metastasis | ER+/PR-/HER2- | Pleural effusion | Letrozole + Goserelin + Abemaciclib; Paclitaxel | ER+/PR-/HER2- |

**Supplementary Table 2: Antibodies used for flow cytometry.**

| Antigen | Fluorophore | Clone | Product reference | Lot number | Manufacturer | Working concentration |
|---|---|---|---|---|---|---|
| ROR1 | APC | REA1051 | 130-118-015 | 5210502199 | Miltenyi Biotec | 1:100 |
| CD3 | PE/Cyanine7 | OKT3 | 317334 | B321207 | BioLegend | 1:400 |
| CD4 | APC/Fire750 | A161A1 | 357426 | B292855 | BioLegend | 1:100 |
| CD8a | PerCP | HIT8a | 300922 | B336677 | BioLegend | 1:50 |
| CD25 | BV421 | BC96 | 302630 | B311394 | BioLegend | 1:50 |
| CD69 | FITC | FN50 | 310904 | B319620 | BioLegend | 1:50 |
| CD31 | FITC | REA730 | 130-110-806 | 5220909327 | Miltenyi Biotec | 1:50 |
